# Modulation by NPYR underlies experience-dependent, sexually dimorphic learning

**DOI:** 10.1101/2023.10.19.563073

**Authors:** Sonu Peedikayil-Kurien, Rizwanul Haque, Asaf Gat, Meital Oren-Suissa

## Abstract

The evolutionary paths taken by each sex within a given species sometimes diverge, resulting in behavioral differences. Given their distinct needs, the mechanism by which each sex learns from a shared experience is still an open question. Here, we reveal sexual dimorphism in learning: *C. elegans* males do not learn to avoid the pathogenic bacteria PA14 as efficiently and rapidly as hermaphrodites. Notably, neuronal activity following pathogen exposure was dimorphic: hermaphrodites generate robust representations, while males, in line with their behavior, exhibit contrasting representations. Transcriptomic and behavioral analysis revealed that the neuropeptide receptor *npr-5*, an ortholog of the mammalian NPY receptor, regulates male learning by modulating neuronal activity. Furthermore, we show the dependency of the males’ decision-making on their sexual status and demonstrate the pivotal role of *npr-5* as a modulator of incoming sensory cues. Taken together, we portray sex-specific plasticity in behavior toward a shared experience by modulating learning.

## Introduction

Learning facilitates the organism’s ability to cope with a changing environment, making it an inherent property of biological neural systems^1^. To adapt to the environment, the organism needs to retain the learning experience and then use the stored information to alter future responses^2^. The synaptic basis of learning and memory has been well established and provides a solid foundation to assess the relationship between experience-dependent learning and survival^3^.

Since sexual selection predicts that males and females are subjected to different selection pressures leading to dimorphisms^4,5^, it stands to reason that the same would occur for an essential ability such as learning. Consistent with this prediction, several sex-shared traits such as perception and learning have been previously demonstrated to be modulated by the genetic sex^6–12^. It has also been proposed that learning can be a driving force for evolutionary change^13^. Thus, sex-specific evolutionary drives might be in tension with decision-making processes when facing an environmental cue^14,15^. While a significant portion of sex differences in behavior could be attributed to the neurochemical pathway of circulating sex hormones, it has been suggested that behavior is also influenced by other factors, such as life history and sex-specific gene expression^16^. This indicates a bi-directional relationship between sexual identity and the environment, with far-reaching consequences on cognitive abilities, including learning and memory. How sexual identity drives molecular and circuit changes toward a shared experience that intersects with learning and survival is still an open question.

The nematode *C. elegans* has been a useful model for the study of learning and memory at the molecular, cellular, neural circuitry, and behavioral levels^17^. Numerous studies have demonstrated that *C. elegans* is capable of non-associative learning and short-term memory, in addition to associative learning, long-term memory, and sensory integration of conflicting cues^18–22^.

The two sexes of *C. elegans* display sexually dimorphic behaviors, such as mechanosensory responses to tail touch, dosage-dependent differences in nociceptive behaviors, and chemoattraction to odors and chemicals. Some of these differences have been mapped to corresponding network changes ^8,23,24^.

Here, we investigated if and how the genetic sex affects context-and experience-dependent behavioral plasticity when learning a shared environmental cue. We harnessed the well-established model of short-term pathogenic aversive conditioning in *C. elegans* ^21,25,26^, where hermaphrodites shift their preference from naïve attraction to aversion upon exposure to the pathogenic bacterium *Pseudomonas aeruginosa* (PA14)^25,27^. We show that males are unable to avoid PA14 as efficiently as hermaphrodites after training. This dimorphic avoidance, however, is not due to differences in the immune responses, but due to layered differences in the sensory and subsequent downstream processing. We observed context-dependent learning when exposing the males to hermaphrodites, which altered their behavior towards PA14. Transcriptomic analyses indicate neuromodulatory regulation, with one candidate, *npr-5*, regulating both the learned as well as context-dependent avoidance by modulating sensory responses. We also unravel the importance of tissue-specific sexual identity and decentralized molecular signaling in regulating male responses. Taken together, our work underscores the diverse levels at which sexual identity modulates dimorphic behavioral responses in each sex towards a shared experience.

### Aversive olfactory learning is sexually dimorphic

To test whether short-term aversive learning is sexually dimorphic, we separated virgin males and hermaphrodites and subjected them to an acute training paradigm of pathogenic bacterial exposure (PA14 or *Escherichia coli* strain OP50 as control) that lasted 6 hours, followed by a choice-based test (Fig. 1a). We found that while hermaphrodites efficiently avoided PA14 at the end of the 6-hour mark, males did not display learned aversion (Fig. 1b-c). A similar dimorphic response was observed when we tested the learning abilities of the sexes against another pathogenic bacterium, *Staphylococcus aureus*^28,29^ (Supplementary Fig. 1a-b). Analogously, when we tested the naïve avoidance by exposing either sex to a lawn of PA14 and scored lawn occupancy after 6-8 hours, males remained within the PA14 lawn while the hermaphrodites’ occupancy was minimal (Supplementary Fig. 1c). We speculated that these observed dimorphisms in short-term learning could arise from differences at checkpoints of information processing within the sexes, such as i) immune responses, ii) integration of sensory information, or iii) a deliberately chosen dimorphic outcome to favor a sex-specific need, in a temporal manner.

**Figure 1.**
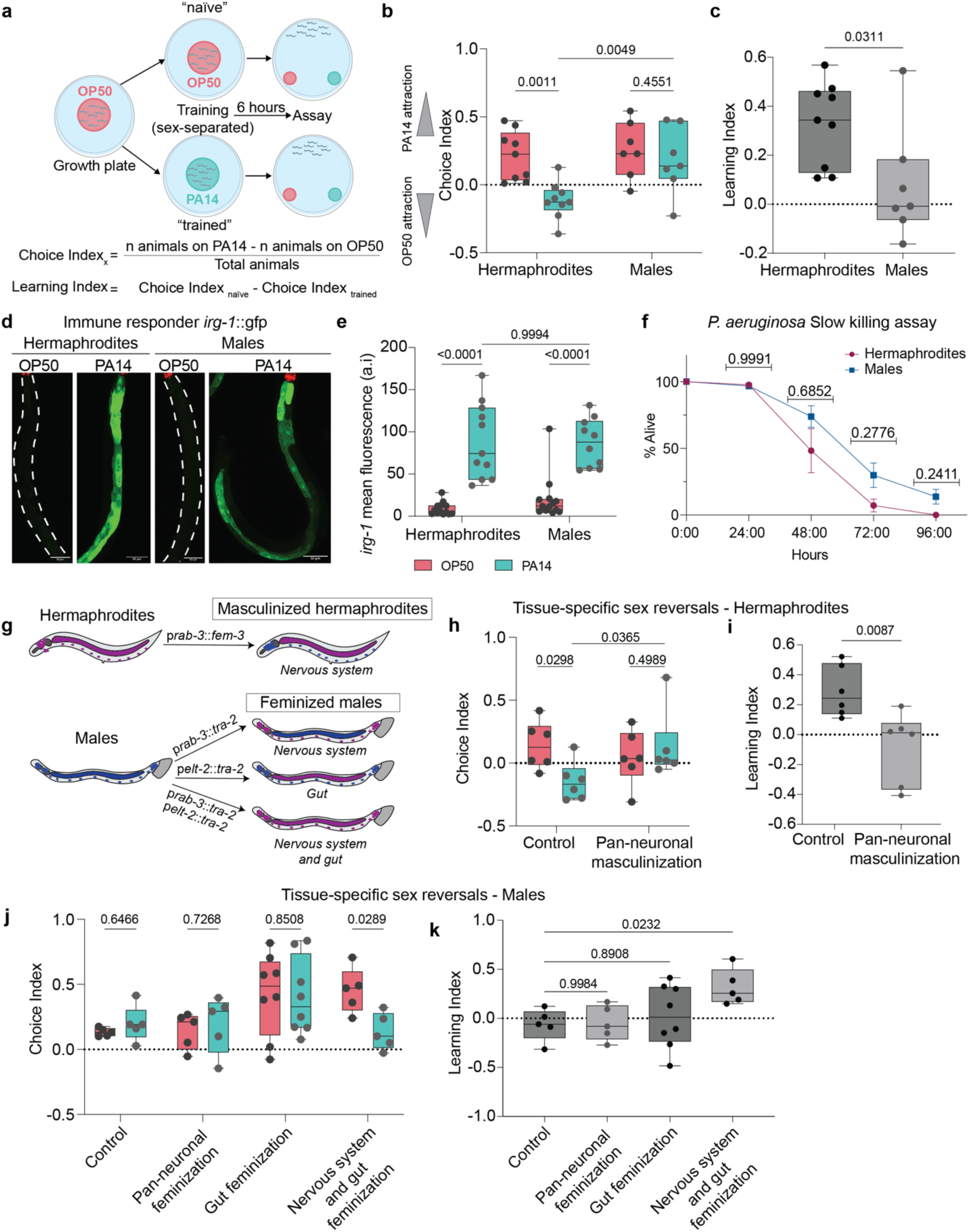
Distinct tissues govern sexually dimorphic short-term learning of PA14 avoidance. **a** Schematic representation of the training paradigm. A group of *C. elegans* was separated into males or hermaphrodites after late L4, exposed to the pathogenic bacterium *Pseudomonas aeruginosa* (PA14, cyan, “trained”) or OP50 (control, magenta, “naïve”) for 6 hours, and then assayed for their choice index against OP50 and PA14 (See Methods). In all figures, n represents the number of animals tested, and N represents the number of biological repeats. **b** The Choice Index of hermaphrodites (OP50, PA14: N = 9) and males (OP50, PA14: N = 7) subjected to the training paradigm in A shows dimorphic responses after training. **c** The learning indices of hermaphrodites (N = 9) and males (N = 7) are dimorphic. **d** Representative confocal micrographs of *irg-1*::GFP hermaphrodites and males trained on OP50 or PA14 after 6 hours of training. Scale bar = 50μm. **e** Quantification of the anterior region of the gut for *irg-1*::GFP in hermaphrodites (OP50: n = 13; PA14: n = 11; n = individual animals) and males (OP50: n = 13; PA14: n = 10; n = individual animals) after exposure to OP50 or PA14. **f** The percentage of live hermaphrodites (N = 6) and males (N = 6) in slow-killing assay. **g** Schematic representation of the tissue sex-reversal experiments (the results of the equivalent germline experiments are presented in Supplementary Fig. 1). **h-i** Choice **(h)** and Learning **(i)** Indices of wild-type (OP50, PA14: N = 6) and pan-neuronally masculinized (p*rab-3*::*fem-3*) hermaphrodites (OP50, PA14: N = 6). **j-k** Choice (**j**) and Learning (**k**) indices of wild-type (control) males (OP50, PA14: N = 5), pan-neuronally feminized (p*rab-3*::*tra-2(ic*)) males (OP50, PA14: N = 5), gut-feminized (p*elt-2*::*tra-2(ic*)) males (OP50, PA14: N = 8), and pan-neuronally and gut feminized (p*rab-3*::*tra-2(ic*);p*elt-2*::*tra-2*) males (OP50, PA14, N = 5). Statistical test used are: in (**b**, **e**, **f**, **h**, **j**), a two-way ANOVA with Šídák’s multiple comparisons test, in (**c**, **i**), a Mann Whitney test. In (**k**), a Brown-Forsythe and Welch’s ANOVA with Dunnett’s T3 multiple comparisons test. Box and Whisker plots with mean and entire data range. P values are indicated in the figure.

To determine whether the immune system is activated in both sexes, we examined candidate genes from multiple pathways activated during bacterial infection using either real-time qPCR analysis or reporter expression levels^30^. The CUB-domain containing gene F08G5.6^31^ and ShK-like toxin gene F49F1.6, mainly activated by the PMK-1 p38 MAPK pathway^32^ were elevated in both sexes after exposure to PA14 (Supplementary Fig. 1d-e). The *pmk-1* pathway-independent immune responder *fshr-1*^33,34^ and its effector, F01D5.5 ^35^, another ShK-like toxin gene, were also elevated in both sexes after training (Supplementary Fig. 1f-g). The *pmk-1* independent gene *irg-3* ^30,35^ was also elevated in both sexes, but to a lesser extent in males (Supplementary Fig. 1h). In addition, we imaged the activation of the infection responder gene *irg-1*, which has been shown to be dependent on *zip-2,* a transcription factor specifically activated upon PA14 exposure and is independent of *pmk-1* ^30^. We found the *irg-1* levels in the anterior gut region of both sexes to be similar (Fig. 1d-e). The similar pathogenic sensitivity was reflected in killing assays conducted to measure PA14 virulence, where both males and hermaphrodites died with similar kinetics (Fig. 1f, Supplementary Fig. 1i). This is in line with the observation that males do indeed avoid PA14, but only after much more extended training periods (Supplementary Fig. 1j-k). Thus, even though the immune response of the males was activated, they were unable to integrate the pathogenicity of PA14 towards learned aversion as rapidly as hermaphrodites.

To evaluate if the avoidance of PA14 is either developmentally timed or inherently dimorphic, we subjected juvenile animals to the same training paradigm as the adult animals (See Methods). We observed that juveniles of both sexes failed to learn and actively preferred PA14 even after training (Supplementary Fig. 1l-n). This suggests that the dimorphic behavior observed in adulthood results from a developmentally mature nervous system^24^.

To characterize the impact of sexual identity on the learning axis, we feminized or masculinized specific tissues in hermaphrodites and males, respectively, and subjected them to the learning paradigm (Fig. 1g). When only the nervous system is sex-reversed, hermaphrodites exhibited a dramatic reduction in their learning ability (Fig. 1h-i). Contrary to our expectations, pan-neuronally feminized males failed to learn (Fig. 1j-k), even though the sex reversal shifted the olfactory preferences of feminized males for certain odors^24^ (Supplementary Fig. 1o), suggesting that the substrate of learned avoidance is restricted mostly to the nervous system in hermaphrodites, but involves additional systems in males. In the absence of a direct link between neuronal regulation and learning in males, we hypothesized that non-neuronal tissues such as the gut and the germline might affect male learning. Indeed, previous studies have shown the importance of the gut in regulating behavior^36,37^, yet feminizing the gut of males was also insufficient to enhance learning in males (Fig. 1j-k). Similar results were obtained when analyzing males with a mutated germline, although germline-mutated hermaphrodites seemed to display dampened avoidance (Supplementary Fig. 1p-q). However, when both the nervous system and the gut were feminized, these males learned to avoid PA14 significantly better than wild-type males (Fig. 1j-k). Thus, the default male drive is orchestrated by several tissues, and their intercommunication is critical for the evaluation and subsequent response to external stimuli.

### Sexually dimorphic neuronal representations of PA14 exposure

We previously demonstrated that while nociceptive sensory neurons in *C. elegans* respond similarly in the two sexes to a noxious cue (glycerol), the network topology diverges, shaping sexually dimorphic behaviors^8^. The neurons regulating PA14 avoidance have been previously mapped^38,39^. The sensory neuron AWC is required for odor-mediated attraction, while AWB is required for odor-mediated repulsion^40–42^. In addition, these neurons have been recently established as important in recognizing the odor bouquet presented by PA14 and for regulating both adult and larval learning^35,38,39^. The dimorphic behavior we observed could then arise from distinct computing of environmental cues (i.e., dimorphic neural states lead to dimorphic behavior). To evaluate this, we used calcium imaging to measure the neuronal activity of sensory- and inter-neurons of naïve and trained hermaphrodites and males and quantified the resulting amplitudes during exposure to a sequence of OP50-PA14-OP50 supernatants (see methods). First, we found higher calcium activity in naïve males, upon presentation of PA14, in the sensory neuron AWC (OFF response) (Fig. 2a-b, i, Supplementary Fig. 2a, c, q) and AWB (ON and OFF response) (Fig. 2c-d, j, Supplementary Fig. 2e, g, r-t), compared to naïve hermaphrodites. Next, we examined the change in response after PA14 training. In both sexes, AWC exhibited no change in OFF response after training, compared to naïve animals (Fig. 2a-b, i, Supplementary Fig. 2a-d, q). However, the repellent odor-sensitive AWB displayed distinct dynamics. While there was no change in the ON response of either sex after training (Fig. 2c-d, Supplementary Fig. 2r-s), the OFF response was higher in hermaphrodites (Fig. 2c-d, j, Supplementary Fig. 2t). Taken together, these results suggest that naïve responses in males are stronger in both AWB and AWC, while hermaphrodites showed stronger trained response in AWB (Fig. 2m).

**Figure 2.**
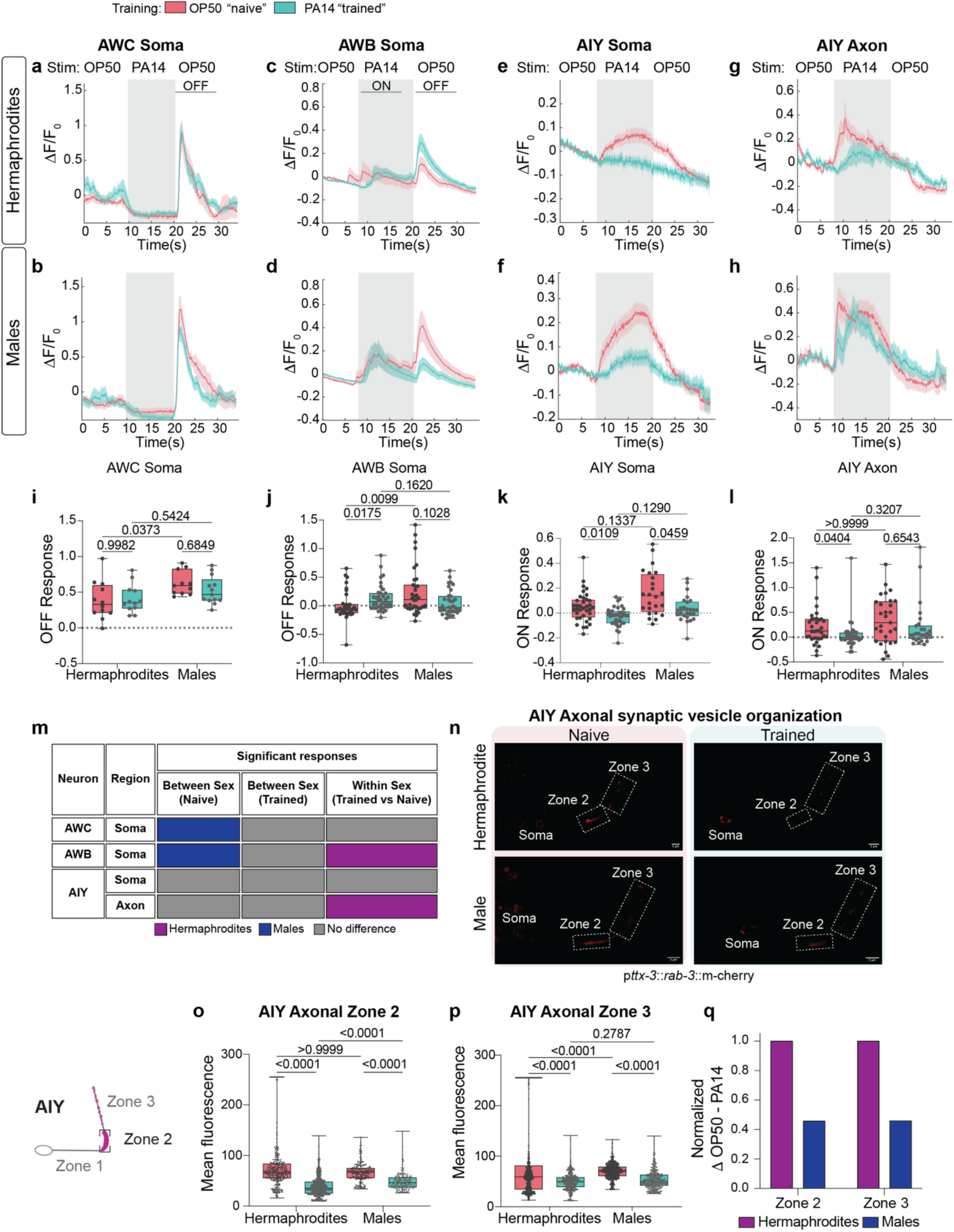
Distinct dimorphic compartmentalized information processing in response to PA14 stimulus. **a-h** From top to bottom, SEM traces of naïve (magenta) and trained (cyan) animals of both sexes in select neurons. The shaded region in the SEM traces corresponds to the exposure window to PA14 supernatant in both naïve and trained animals. OP50 supernatant was used as the buffer (See Methods). **a** AWC calcium levels in trained (n = 12) and naïve (n = 12) hermaphrodites. **b** AWC calcium levels in trained (n = 13) and naïve (n = 11) males. **c** AWB calcium levels in trained (n = 36) and naïve (n = 34) hermaphrodites. **d** AWB calcium levels in trained (n = 35) and naïve (n = 35) males. **e** AIY soma calcium levels in trained (n = 36) and naïve (n = 34) hermaphrodites. **f** AIY soma calcium levels in trained (n = 35) and naïve (n = 35) males. **g** AIY axonal calcium levels in trained (n = 35) and naïve (n = 35) hermaphrodites. **h** AIY axonal calcium levels in trained (n = 27) and naïve (n = 29) males. **i** Quantification of OFF response of AWC. **j** Quantification of OFF response of AWB. **k** Quantification of ON response of AIY. **l** Quantification of ON response of AIY Axon. **m** Table summarizing the neuronal responses observed. Purple, significant response in hermaphrodites; Blue, Significant response in males; Grey, no difference between the sexes. The sensory neurons AWC and AWB showed higher calcium responses in the naïve males compared to naïve hermaphrodites. After training, hermaphrodites exhibit dimorphic OFF responses in AWB, and ON response in AIY axon. **n** Representative confocal micrographs of p*ttx-3*::*rab-3*::mCherry in naïve and trained hermaphrodites and males showing expression in distinct zones of AIY, namely, a synapse-devoid Zone 1 and synapse-rich Zones 2 and 3. Scale bar = 12μm. **o-p** Synaptic RAB-3 mCherry fluorescence in AIY Zone 2 and Zone 3 axonal regions, respectively. Each dot represents a puncta from 26-47 animals per group. **q** Normalized delta between naïve and trained RAB-3 expression of both sexes in Zone 2 and Zone 3. Hermaphrodites in magenta, and Males in blue. Data is normalized to hermaphrodites. In (**a-h**) change in fluorescence (ΔF) was normalized to the average GCaMP signal in 10 seconds before stimulus start (F_0_). ON response was defined as the difference in average ΔF/ F_0_ during 10 seconds after stimulus start to 5 seconds before stimulus start. OFF response was calculated as the difference in average ΔF/ F_0_ during 10 seconds after stimulus removal to 5 seconds before stimulus start. In (**i-l**, **o-p**), the statistical test used is Kruskal-Wallis test with Dunn’s multiple comparisons test was performed. Box and Whisker plots with mean and entire data range. P values are indicated in the figure.

The sensory information from AWC and AWB is passed on to a downstream interneuron AIY, which has been implicated as an information checkpoint that regulates memory retrieval ^39,43–45^ as well as hermaphrodite training and forgetting states of PA14 aversive learning ^26,38^. We observed a reduction in calcium activity after training at the soma of the AIY neuron in both sexes (Fig. 2e-f, k, Supplementary Fig. 2i-l, u). Yet, within the axonal regions of AIY, we observed a decrease in calcium activity after training only in hermaphrodites but not in males (Fig. 2g-h, l, Supplementary Fig. 2x). Interestingly, the latency of calcium responses in AIY soma (Supplementary Fig. 2v) and axonal regions (Supplementary Fig. 2y) of trained males was smaller when compared to trained hermaphrodites, while the time the neuron was active was unchanged in all the groups examined (Supplementary Fig. 2w, z). Taken together, our results show an increased sensory response to PA14 in AWB, and a subsequent decrease in the interneuron AIY only in hermaphrodites. However, the neuronal response landscape was dimorphic with naïve male sensory responses higher and the interneuron AIY displaying compartmentalized (soma vs axonal) dimorphic responses (Fig. 2m).

To determine whether the observed dimorphisms in neuronal activity also extend to the synaptic vesicle organization of AIY, i.e., a readout of the learning process, we used a fluorescent readout of RAB-3, which regulates docking of synaptic vesicles^46–48^. The axonal regions of AIY can be distinguished according to their respective synaptic partners as Zone 2 and 3 (Fig. 2n) ^48^. We quantified the mCherry expression levels of RAB-3 after training with PA14 and observed a non-dimorphic decrease in RAB-3 levels in both Zones 2 and 3 (Fig. 2o-p), suggesting that the sensory stimulus is processed in both sexes. However, we noticed that the extent of difference in RAB-3 expression between the naïve and trained males is lower than that of hermaphrodites (Fig. 2q), suggesting that dimorphic perceptive information is being communicated downstream.

### Dimorphic molecular landscape of naïve and trained animals

To characterize the molecular landscape of short-term dimorphic learning, we performed whole-animal transcriptomics on sex-separated naïve and trained animals of both sexes (4 groups with 4 biological repeats), and the resultant RNA was sequenced using a bulk optimized protocol of the single-cell MAR-seq protocol^49^. This protocol has been previously used to establish the molecular profile of both sexes in all developmental stages^50^. To visualize the relationship between training and sexual identity across all experimental groups, we performed a principal component analysis (PCA) on the ∼ 12,000 identified genes. The PCA revealed a clear separation by sex (Fig. 3a), validated by the dimorphic expression of *lsy-27, tbx-9* and *tra-2* (higher in hermaphrodites^50,51^) and *mab-3, pkd-2* and *ins-39* (higher in males^50,52^) (Supplementary Fig. 3a). The training status of the animals was also separated in the PCA (Fig. 3b). We found 869 and 846 differentially expressed genes in hermaphrodites and males, respectively, after training (p value < 0.05; Supplementary Data 1). Unlike the behavioral results, where both naïve and trained males displayed similar attraction towards PA14 (Fig. 1b-c), their molecular landscape differed and reflected the training experience (Fig. 3b). Volcano plots and functional GO term analysis revealed similar changes in immune regulation, fatty acid metabolism, and other stress-related pathways in both sexes, an expected outcome given the gustatory and pathogenic nature of the training bacterium (Fig. 3c-d, Supplementary Fig. 3c, Supplementary Data 1, Supplementary Data 2). Of the entire transcriptomics dataset, 70 genes were previously reported to be involved in PA14-related behaviors (Supplementary Data 2)^26,27,37,39,53–64^. Consistent with our survival assays, most of the genes implicated in PA14 survival, such as *daf-16* and *daf-7*, were non-dimorphic. Genes involved in training and forgetting, such as *gst-38* and *gst-26* ^26^, were similarly regulated in trained animals of both sexes (Supplementary Fig. 3d, Supplementary Data 2). qPCR analysis of metabolic gene *acdh-1* ^65^, implicated in PA14 survival, was reduced non-dimorphically after PA14 exposure as indicated by our transcriptomics dataset (Supplementary Fig. 3b, d).

**Figure 3.**
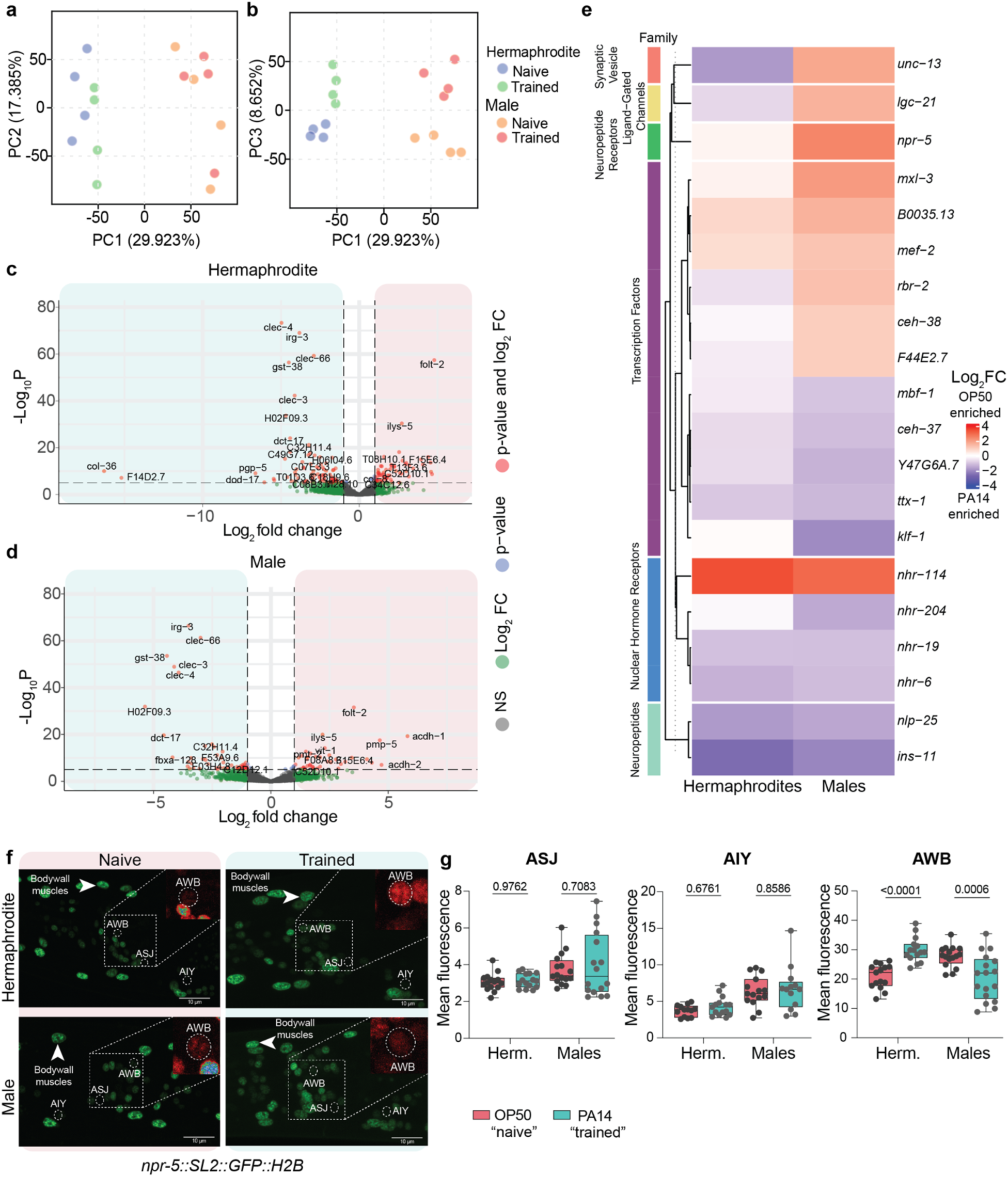
Sexually dimorphic learning triggers sex-specific changes in NPYR expression. **a-b** Principle Component Analysis (PCA) of the transcriptomes of hermaphrodites and males (n = 4 independent samples from 600-800 animals/replicate) after training, ordered according to sex (**a**) and training status (**b**), showing the validity of the data in separating the sexes and training states by transcriptomics. **c-d** Volcano plots of naïve and trained hermaphrodites (**c**) and males (**d**), depicting similar changes in transcripts **e** Heatmap of log_2_ fold-change of neuron-relevant genes from naïve vs. trained males (FDR < 0.05) and hermaphrodites. Red indicates enrichment in naïve animals (OP50), while blue indicates enrichment in trained animals (PA14). See also Supplemental tables 1-2. **f** Representative confocal micrographs of *npr-5::SL2::GFP::H2B* CRISPR knock-in in naïve and trained hermaphrodites and males showing expression in body wall muscles and sensory cells. The inset shows the heatmap expression of *npr-5::SL2::GFP::H2B* in the selected region with AWB and ASJ. Scale bar = 10μm. **g** Quantification of mean fluorescent signals of *npr-*5 in ASJ, AIY and AWB, showing dimorphic expression changes in AWB after training (ASJ (hermaphrodites: OP50, PA14 n =15; males: OP50, PA14 n =16); AIY (hermaphrodites: OP50 n =13, PA14 n =15; males: OP50 n =15, PA14 n =13); AWB (hermaphrodites: OP50, PA14 n =15; males: OP50 n =16, PA14 n =17). In **(g)**, statistical test used is two-way ANOVA with Šídák’s multiple comparisons test. Box and Whisker plots with mean and entire data range. P values are indicated in the figure.

To identify transient changes that regulate male learning, we focused on differentially expressed neuro-specific gene classes, such as receptors and neuropeptide families (Fig. 3e, Supplementary Data 2). We uncovered changes in transcription levels for several transcription factors. The homeodomain *ceh-37*, which specifies the identity of the sensory neuron AWB^66^ and is involved in regulating immunity from the gut^67^ was upregulated in both sexes (Fig. 3e). *mxl-3*, involved in lipolysis and expressed in sensory neurons including ASE, AWC and AWB^68,69^ was downregulated in males, while the H3K4 demethylase *rbr-2*, involved in axon guidance and transmission of stress resistance across generations^70,71^ was dimorphically regulated (Fig. 3e). Interestingly, the levels of neuropeptide *ins-11*^61^, a negative regulator of PA14 avoidance, were increased in trained animals of both sexes (Fig. 3e), as confirmed by qPCR analysis (Supplementary Fig. 3b, d). *unc-13*, a gene involved in synaptic vesicle clustering, was observed to be dimorphically regulated after training in the sexes, along with the ligand-gated channel *lgc-21* and the neuropeptide receptor *npr-5*. qPCR analysis confirmed the dimorphic expression of *npr-5,* which is down-regulated in males after training (Fig. 3e, Supplementary Fig. 3b), but not of *unc-13* (Supplementary Fig. 3b). Overall, our transcriptomic data suggest that while both sexes respond to PA14 exposure with a similar molecular signature, additional modulatory mechanisms may have evolved to regulate the observed male behavior.

### Sexually dimorphic modulation of the NPY receptor expression

Recent evidence suggests the importance of integration between the nervous and the immune systems in generating appropriate responses towards pathogenic bacterium^72^. Of note, neuropeptides and their receptors play a major role in PA14 avoidance behavior^73^. One such example is the G-protein coupled receptor *npr-1,* which has been shown to inhibit AQR, PQR, and URX neurons to confer resistance to PA14^74,75^. Given the importance of the neuropeptidergic system in modulating diverse behaviors, including learning^76^, the neuropeptide receptor *npr-5* emerged as a likely candidate. Notably, NPR-5 is an ortholog of the mammalian NPY receptor, a key regulator of various behaviors, including learning and memory^77–81^. In *C. elegans*, *npr-5* is involved in regulating foraging and fatty acid metabolism in hermaphrodites^82,83^. In addition, *npr-5* was shown to modulate feeding by regulating the secretion of serotonin from ADF neurons in an NMDAR-dependent manner^84^, both of which have independently been implicated in PA14 aversive learning ^27,85^. Thus, we sought to examine its role in male learning behavior.

To determine the change in expression post-training, we endogenously tagged *npr-5* using CRISPR-Cas9 gene-editing and quantified its expression levels in head sensory neurons, previously described to regulate behavior^82,83^ (Fig. 3f-g). Consistent with previous studies^82^, *npr-5* was observed in body wall muscles and multiple neurons, including sensory neurons (Fig. 3f). After training, *npr-5* levels rose in hermaphrodite AWB neurons but were reduced in male AWB neurons (Fig. 3g). Intriguingly, this change in *npr-5* expression resembles the changes in activity levels of AWB (Fig. 2c-d). Although ASJ has been previously described to be involved in processing PA14 experience^37,64,86^, and AIY displayed dimorphic response (Fig 2g, h, l), no significant changes were observed in *npr-5* expression after training in ASJ and AIY (Fig. 3g). These results support the transcriptomic data, suggesting *npr-5* is down-regulated in males upon PA14 exposure (Fig. 3e), pointing to a critical role of *npr-5* in modulating learning, perhaps at the sensory level.

### Neuropeptidergic modulation of sex-specific learning

To evaluate the extent of neuropeptidergic regulation in male-specific PA14 aversive learning, we subjected *npr-5* mutants to the training paradigm. Strikingly, males with a partial or full *npr-5* deletion acquired the ability to learn quickly in a similar fashion as wild-type hermaphrodites (Fig. 4a-d). Although mutant *npr-5* hermaphrodites still avoided PA14, no significant changes were observed between the naïve and trained responses (Fig. 4a, c). Previous studies^82^, along with our calcium imaging (Fig. 2c-d, j) and *npr-5* expression results (Fig. 3f-g) suggest a sensory-specific site-of-action of *npr-5*. Therefore, we rescued *npr-5* expression specifically in AWB (using the *str-1* promoter) and in ciliated neurons (using *tax-4* promoter) in an *npr-5* mutant background. While both manipulations sufficed to impede the decision-making ability as indicated by the choice index (Fig. 4a), reverting the animal back to the wild-type male phenotype, the *tax-4* rescue strain displayed a stronger phenotype in the learning index (Fig. 4b). Thus, *npr-5* is required for suppressing learned avoidance, and its activity, possibly in AWB and other sensory neurons, suggests a sensory gating mechanism in males, in line with the observations made in the neuronal activity of trained males (Fig. 2).

**Figure 4.**
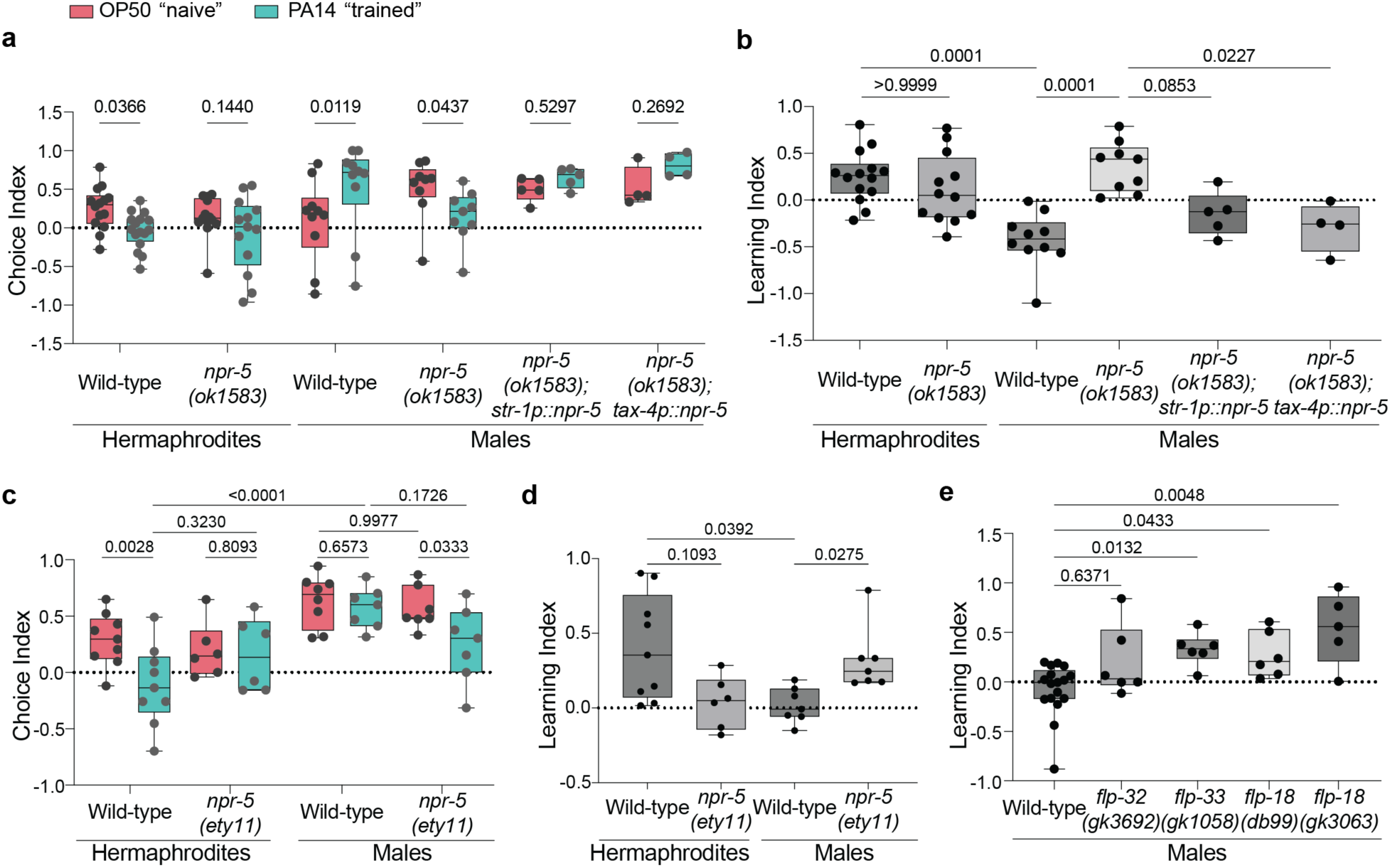
NPR-5 regulates male-specific behavior. **a-b** Choice (**a**) and Learning (**b**) Indices of naïve (magenta) and trained (cyan) wild-type and *npr-5* mutant hermaphrodites and males and male rescue strains where *npr-5* cDNA was expressed either under *str-1* promoter (AWB specific expression ^88^) or *tax-4* promoter (expression in ciliated neurons ^89^); data shows sex-specific behavioral effects in males (Choice Index: wild-type hermaphrodites: OP50 (N = 12), PA14 (N = 13); *npr-5* hermaphrodites: OP50 (N =10), PA14 (N = 12); wild-type males: OP50, PA14 (N = 10); *npr-5* males: OP50, PA14 (N = 9); p*str-1*::*npr-5*cDNA; *npr-5* males: OP50, PA14 (N = 5); p*tax-4*::*npr-5*cDNA;*npr-5* males: OP50, PA14 (N = 4)). Learning Index: wild-type hermaphrodites: (N = 12); *npr-5* hermaphrodites: (N = 11); wild-type males: (N = 10); *npr-5* males: (N = 9); p*str-1*::*npr-5*cDNA;*npr-5* males: (N = 5); p*tax-4*::*npr-5*cDNA;*npr-5* males: (N = 4)). **c-d** Choice (**c**) and Learning (**d**) indices of naive (magenta) and trained (cyan) wild-type and *npr-5* CRISPR mutant hermaphrodites and males, showing sex-specific behavioral effects in males (hermaphrodites: OP50, PA14, Learning Index (n = 9); *npr-5* hermaphrodites: OP50, PA14, Learning Index (N = 6); males: OP50, PA14, (N = 8), Learning index (N = 7); *npr-5* males: OP50, PA14, Learning Index (N = 7)). **e** Learning indices of wild-type (N = 17), *flp-32* (*gk3692*) mutant, *flp-33* (*gk1058*) mutant, *flp-18* (*db99*) mutant and *flp-18 (gk3063)* males (N = 6), showing peptidergic regulation of male learning. Statistical test used are: in (**a, c**), a two-way ANOVA with Tukey’s multiple comparisons test, in **(b, d, e)**, a Kruskal-Wallis test with Dunn’s multiple comparisons test. Box and Whisker plots with mean and entire data range. P values are indicated in the figure.

Previous studies have also established the link between various neuropeptides and the regulation of behavioral states^86,87^. Given the dimorphic neuropeptidergic landscape and the decentralized learning processing we observed in males (Fig. 1j-k), we postulated that the learning behavior in males might be regulated by multiple signaling molecules in a molecular network (Supplementary Fig. 4a). Indeed, when we screened select candidate neuropeptides using mutants or RNAi, we observed that several neuropeptides such as *flp-33*, *flp-3* and *flp-18*, a cognate ligand of *npr-5,* were involved (Fig. 4e, Supplementary Fig. 4b-c). The regulation of male PA14 aversive learning by *npr-5* and multiple neuropeptides demonstrates that male learning is an amalgamation of complex internal processing.

### *npr-5* functions in AWB to modulate PA14 responses

Based on its sensory responses to PA14 (Fig. 2c-d, j), the observed decrease in *npr-5* expression following training (Fig. 3g), and the change in behavior upon rescue (Fig. 4a), AWB is a likely target of *npr-5* action. We, therefore, measured AWB activity in *npr-5* mutants and observed a striking change in calcium response patterns, namely, a heightened OFF response (Fig. 5a-d, Supplementary Fig. 5a-b), compared to wild-type animals (Fig. 2c-d). In addition, only *npr-5* mutant male AWB neurons exhibited a stronger response to the removal of the PA14 stimulus after training (Fig. 5c-d), similar to wild-type hermaphrodites (Fig. 2c, j). The *npr-5* hermaphrodites showed a decrease in both ON and OFF responses after training (Fig. 5a, c, Supplementary Fig. 5a, c-d). Intriguingly, when stimulated with diluted 2-nonanone, an aversive odor recognized by AWB^90^, wild-type animals displayed a non-dimorphic response (Supplementary Fig. 5e, g-h). However, *npr-5* mutants showed higher activation in both sexes when compared to their wild-type counterparts, with *npr-5* males displaying dimorphic heightened activation (Supplementary Fig. 5e-h). Together, our results support the dimorphic inhibitory role of *npr-5* in the proper integration of PA14 signals.

**Figure 5.**
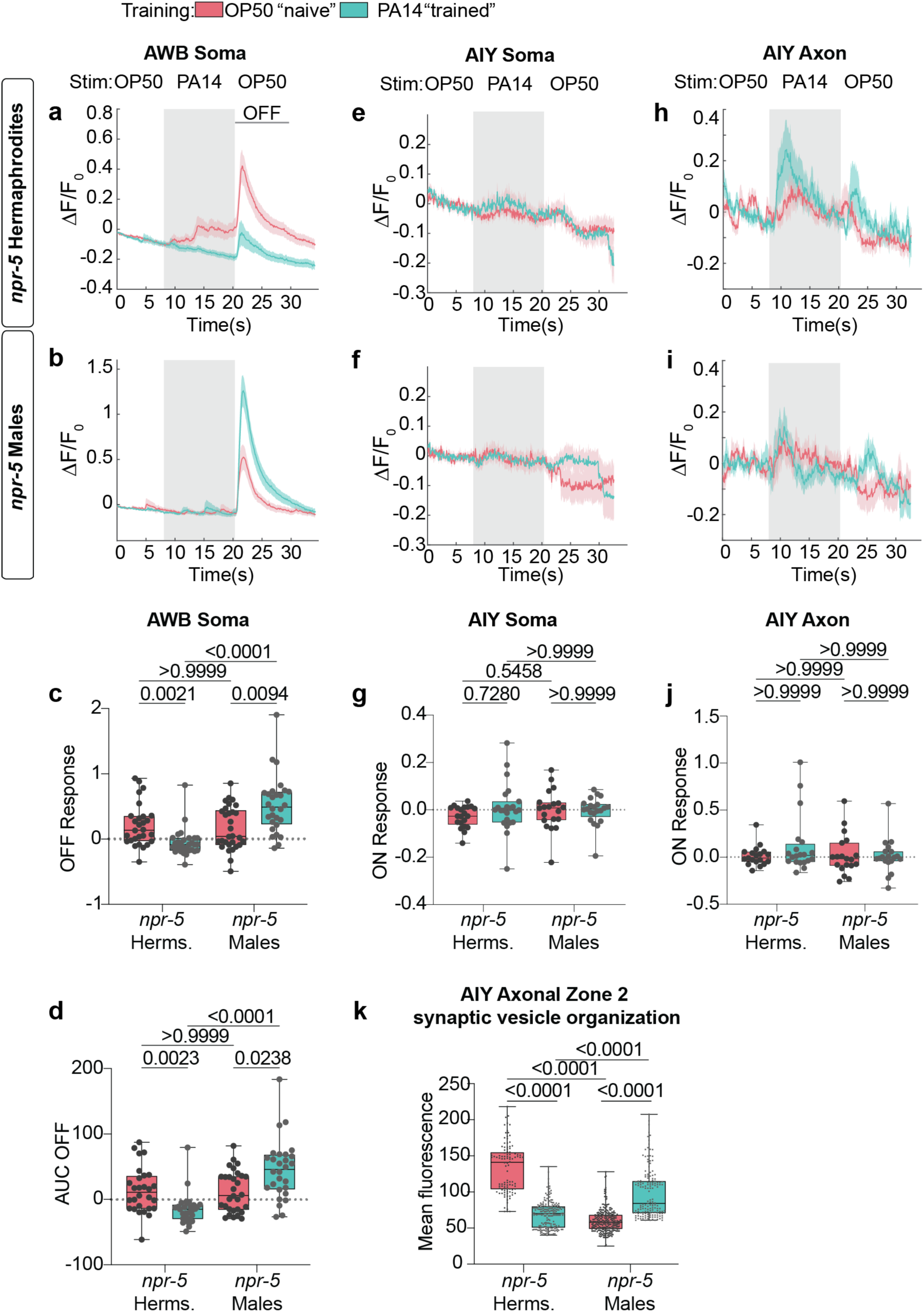
NPR-5 modulates state and sex-specific neuronal activity. **a-b, e-f, h-i** From top to bottom, SEM traces of GCaMP signals of select neurons of *npr-5* hermaphrodites and males in naïve (magenta) and trained (cyan) animals. The shaded region corresponds to the exposure window of PA14 supernatant. OP50 supernatant was used as the buffer (See Methods). AWB soma calcium response of naïve and trained **(a)** *npr-5* mutant hermaphrodites (OP50 n = 28, PA14 n = 29) and **(b)** *npr-5* mutant males to PA14 supernatant (OP50 n = 30, PA14 n = 28). **c-d** Quantification of **c** the OFF response and **d** AUC of OFF response of AWB. **e** AIY soma calcium levels in naïve (n = 21) and trained (n = 23) *npr-5* mutant hermaphrodites. **f** AIY soma calcium levels in naïve (n = 20) and trained (n = 21) *npr-5* mutant males. **g** Quantification of the ON response of AIY soma. **h** AIY axon calcium levels in naïve (n = 20) and trained (n = 19) *npr-5* mutant hermaphrodites. **i** AIY axon calcium levels in naïve (n = 19) and trained (n = 20) *npr-5* mutant males. **j** Quantification of the ON response of AIY axon. **k** Mean quantification of synaptic *rab-3::mCherry* fluorescence in naïve and trained *npr-5* mutant hermaphrodite and male AIY axon. Each dot represents a puncta from n = 10-17 animals per group. In **(a-b, e-f, h-i)** change in fluorescence was calculated as for Figure 2 (see Methods). In **(c-d, g, j, k)**, statistical test used is Kruskal-Wallis test with Dunn’s multiple comparisons test was performed. Box and Whisker plots with mean and entire data range. P values are indicated in the figure.

Even though we observe no change in *npr-5* expression in AIY (Fig. 3g), we found a stark decline in the AIY soma calcium activity in only naïve, but not trained, *npr-5* mutants (Fig. 5e-g, Supplementary Fig. 6a-b, e) compared to wild-type animals (Fig. 2e, f, k, Supplementary Fig. 2i-l), suggesting that *npr-5* is involved in the recognition and processing of PA14 in the naïve state. Similar observations were made in the axonal responses of AIY to PA14 odor in naïve mutant hermaphrodites and males (Fig. 5h-j, Supplementary Fig. 6c-d, h). In contrast to wild-type males, which retained high activation after training (Fig. 2h, l), mutant males showed reduced activation (Fig. 5h-j, Supplementary Fig. 6c-d, h). The lack of change in responses between the naïve and trained states is also reflected in the latency and the activation time of AIY in mutant animals (Supplementary Fig. 6f-g, i-j). Taken together, these results suggest that *npr-5* plays a crucial role in the integration of PA14 naïve cues in both sexes, but also the trained responses of males. This is highlighted also at the synaptic level where the RAB-3 expression of *npr-5* mutant males displayed an increase in mean fluorescence after training (Fig. 5k), unlike wild-type animals (Fig. 2o) and *npr-5* mutant hermaphrodites (Fig. 5k). Thus, exposure to PA14 is processed dimorphically between the sexes, altered partly through modulation by *npr-5* in males.

### NPR-5 overexpression curbs state-specific male behavior

Our pathogen-sensitivity and calcium-imaging results suggest that while males possess the ability to recognize and potentially process the PA14 experience (Fig. 1c-d, Fig. 2), learning is repressed. *C. elegans* males will abandon food in search of mates^24^ and switch from avoidance to attraction of aversive stimuli if hermaphrodites have been experienced^18,91^. Thus, we hypothesized that perhaps repression in our paradigm is a result of the inherent evolutionary drive for mate search. Therefore, diminishing males’ sexual drive could lead to PA14 avoidance. To assess this, we trained animals with the other sex (“sex-exposed”) or without (“virgin”) in the presence of either OP50 or PA14 (Fig. 6a) and checked their choice indices. Sexual conditioning would predict that in the presence of hermaphrodites, the males would be highly attracted to the coupled stimulus, as evidenced by the retention of attraction to OP50 by the naïve hermaphrodite-exposed males (Fig. 6b). Surprisingly, we observed that, as opposed to virgin-trained males, which retain their attraction to PA14, trained hermaphrodite-exposed males show decreased attraction to PA14 (Fig. 6c). Our results suggest that reducing the males’ sexual or mate-searching drive might allow the trained decision-making process to flow uninterrupted. To assess the potential role of *npr-5* in regulating the integration of the internal state into the learning circuit, we overexpressed *npr-5* in sensory neurons of wild-type males to simulate a perpetual ‘naïve’ state (Fig. 6b). These males failed to avoid PA14, even after training, regardless of their sexual status (Fig. 6c). Taken together, these results suggest that sex-specific needs act as a regulator of male learning, dictated and processed by the inherent neuromodulatory state, where *npr-5* in the sensory neurons of males halts potential learning by modulating the diverse variables that affect male decision-making (Supplementary Fig. 7).

**Figure 6.**
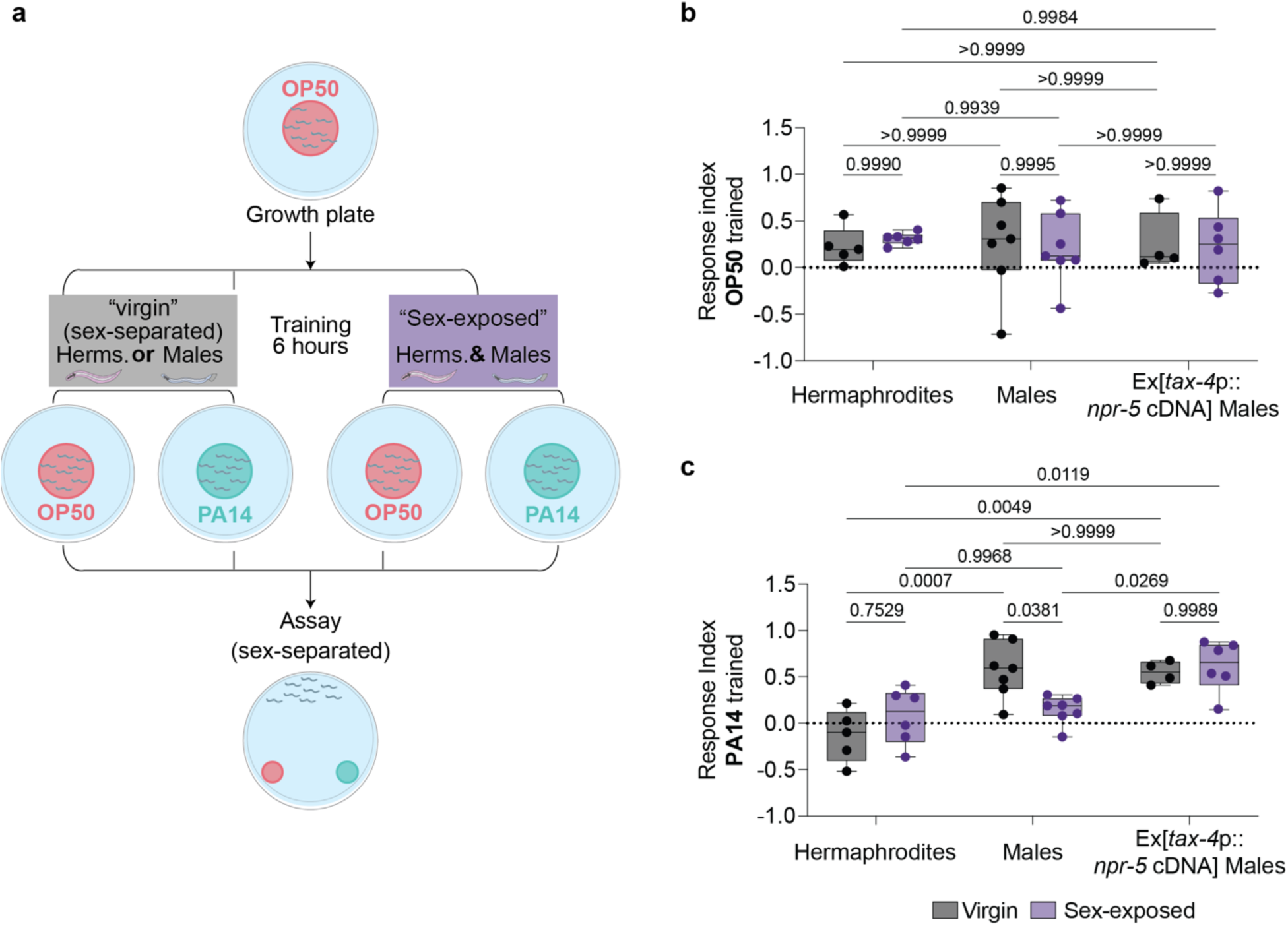
The sexual status determines the behavioral outcomes. **a** Schematic representation of the conditioning training paradigm. A group of *C. elegans* was separated into males or hermaphrodites after late L4, exposed to OP50 (magenta) or PA14 (cyan) for 6 hours either separately (“virgin”) or both sexes together (“sex-exposed”), and then assayed for their choice index against OP50 and PA14 (See Methods). **b** Choice Index of OP50-exposed virgin and sex-exposed wild-type hermaphrodites (virgin N = 5, sex-exposed N = 6), wild-type males (virgin, sex-exposed N = 7) and *npr-5*-overexpression mutant (p*tax-4*::*npr-5*cDNA) males (virgin N = 4, sex-exposed N = 6). **c** Choice Index of PA14-exposed virgin and sex-exposed wild-type hermaphrodites (virgin N = 5, mated N = 5), wild-type males (virgin N = 7, mated N = 7) and *npr-5*-overexpression males (p*tax-4*::*npr-5*cDNA virgin N = 4, mated N = 6). N refers to number of independent assays with each dot representing a single population assay of 50-200 animals/assay. In (**b**, **c**), statistical test used is two-way ANOVA with Tukey’s multiple comparisons test. Box and Whisker plots with mean and entire data range. P values are indicated in the figure.

## Discussion

Sex-specific behaviors have long been recognized as both products and drivers of sexual selection, however learning, a sex-shared trait, would naïvely be expected to withstand the forces of sexual selection, as it is crucial to an organism’s survival. Notably, a recent study demonstrated that in a limited information paradigm, female rats outperform males^92^. These findings underscore the importance of evaluating learning within the context of survival, where information is rapid, limited, and temporal. In our study, we investigated the learning performance of both *C. elegans* sexes in a multi-factorial, ethologically relevant paradigm, specifically focusing on pathogen avoidance. Our results reveal an intriguing behavioral dimorphism: Males do not learn to avoid the pathogenic bacteria PA14 as efficiently and rapidly as hermaphrodites.

The exposure to PA14 generates a neural representation of that experience^26,38,39^. Indeed, we observe that two major olfactory neurons, AWC and AWB, and a cholinergic interneuron AIY, reflect that training experience. Consistent with our notion that males can process the training experience, we observe that the soma of AIY reduce its responses to PA14 supernatant, in a manner similar to that of hermaphrodites, while there was no change in AWC after training in either of the sexes. AWB displayed a biphasic response, with naïve males showing higher ON response than naïve hermaphrodites and remained unaffected by training, while the OFF responses of AWB were sexually dimorphic after training. The ON response in AWB could stem from a possible integration mechanism of odors of both OP50 and PA14, a mechanism previously described by Yang et al^90^ or it is a response to the removal of OP50^38^. The OFF response of AWB increased in hermaphrodites but reduced in males. Parallelly, the axonal regions of male AIY remain unaffected but that of hermaphrodites showed a reduction analogous with their AIY somas. Although compartmentalization of information processing has been shown before^93^, we uncover a possible sexually dimorphic component. The neuropeptide receptor *npr-5*, maintains these neural states as its mutation caused a switch in male AWB OFF responses, which heightened after training, and a complete silencing of naïve AIY responses. These point to the existence of a sensory gating mechanism in the males where information is diverted, partly through *npr-5*, so as to continue choosing PA14 over OP50 until the repression is alleviated.

Thus, in our investigation of the underlying molecular mechanisms that drive these behavioral disparities, we identified several notable divergences in how males and hermaphrodites process shared information during the learning process (Supplementary Fig. 7). First, learning in hermaphrodites appears to be contingent on their nervous system, whereas males require the combined processing of both their nervous system and gut to make critical choices between benign and pathogenic bacteria. This indicates the existence of a complex regulatory mechanism in males that overrides the immediate threat of infection, a finding in line with recent evidence highlighting the importance of the gut-brain axis in behavior regulation^37,94^. Our study further suggests the presence of a sexually dimorphic component. A potential explanation is the dispersed network of neuropeptides we uncovered that could partially function from the gut. Second, our investigation of neuronal correlates of learning revealed distinct sex-specific activation levels and responses to PA14 stimuli in the sensory neurons and interneurons examined. This observation adds another facet to the developing proposal of the sex differences in synaptic mechanisms that underlie plasticity, learning, and memory^7^. Sex hormones have been shown to play a role in learning^95,96^. Importantly, our study highlights the role genetic sex plays in neuronal function independent of sex hormones. Lastly, the MAR-seq analysis of naïve and trained animals points to transcriptomic differences between males and hermaphrodites, particularly within the underlying neuromodulatory network that potentially contributes to the observed behavioral differences. Similar findings have been reported in butterflies, where a genomic analysis revealed a tissue and gene family-specific bias between the sexes that could explain learning dimorphisms^97^. Recent evidence highlights the important role of neuromodulators in recognizing and avoiding PA14^54,58^. Consistent with this, we demonstrate that the sensory modulation of NPR-5 expression is not only crucial for male behavior but also for the maintenance of neuronal calcium response patterns. Further studies are required to examine this effect in other neurons as well as its universality in multiple behavioral contexts.

A key question arises regarding the potential influence of evolutionary pressures. Our data suggest that while males do eventually learn to avoid PA14, they might exhibit short-term repression as a behavioral adaptation. Drawing a parallel to models of foraging animals that exhibit overharvesting or over-staying^98^, it may be evolutionarily beneficial for *C. elegans* males to stay in a patch of pathogenic bacterium, despite immunological activation, to maximize the chances of encountering incoming hermaphrodites. Supporting this notion is the observed reduction in attraction towards PA14 in mated males. One interpretation of this finding is that by removing the need to search for mates, the mating pressure is reduced in males, thus allowing them to activate the otherwise gated behavior. This observation is further supported by recent evidence of increased mating by hermaphrodites previously exposed to pathogenic bacterium^99^. It is tempting to speculate that these two pathways could have co-evolved to maximize organismal fitness under dire conditions. In such a scenario, the overall evolutionary demand of the species is fulfilled, albeit in a complex sex-specific manner. A fascinating example is the lower participation of the male bumblebee in cognitive tasks^100^, even though they share comparable learning abilities with sterile worker females^101^, suggesting that nuanced sex-specific roles alter the perceived learning objective for each sex. Therefore, considering the sex differences in decision-making behavior across organisms, it stands to reason that sexual selection impinges on learning behaviors to favor sex-specific outcomes. By assessing the influence of sexual identity at key levels involved in learning, such as molecular and neuronal circuits, we emphasize the importance of considering sex differences in learning and memory disorders.

## Methods

### Strains and experimental design

Nematodes were cultivated on regular NGM plates on *Escherichia coli* strain OP50 at room temperature unless otherwise mentioned. *him-5*(e1490) were used as a source for wild-type males. All transgenic and bacterial strains are listed in Supplementary Data 3.

### PA14 aversive learning

Animals were age-synchronized by bleaching, unless otherwise specified, and were grown at room temperature on OP50 plates until they were late L4, sex-separated into males and females for behavioral assays as young adults. The training and assay plates were slightly modified from ^27^. Briefly, OP50 and PA14 were streaked from glycerol stocks onto LB plates and incubated at 37°C overnight. They were then moved to 4°C and used for up to a week. To prepare OP50 or PA14 cultures used in the training and assay plates, a small amount of bacteria from streaked plates were incubated in 15 ml tubes containing 5 ml LB at 37°C overnight at 250 rpm. After which, the specified amount of overnight culture (200 μl bacteria for the training plates and 10 μl for the assay plates) was plated onto dried NGM plates and incubated at 37°C overnight and then at 26°C for 24 hours. Before being used for behavioral assays, the plates were acclimatized to room temperature (22-23°C). Care was taken to avoid overly dried or contaminated plates. Before each experiment, a test batch of training and assay plates were verified for behavioral consistency using young adult hermaphrodites, failure of which resulted in fresh streaks from a glycerol stock.

For the training, sex-separated animals were washed using distilled water or M9 and introduced into training plates for 6 hours at room temperature unless otherwise mentioned. Excess liquid was dried with a Kim-wipe. For the choice assay, the worms were washed off the training plates and washed twice in distilled water or M9. Then, they were placed on the choice assay plates and dried with a Kim-wipe. Once dried, they were left at RT for an hour before being scored manually.

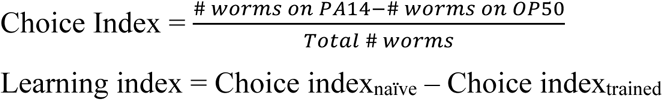

To prepare the worms, L1-arrested worms were grown on regular NGM plates until they were L3, approximately 28 hours. For the training, L3 worms were washed and put on either OP50 (control) or PA14 training plates and then incubated at RT for 6 hours. After which, they were washed off the plates and washed twice in distilled water or M9. Then, they were placed on choice assay plates and dried with a Kim-wipe. Once dried, they were left at RT for an hour and then, using a sharp surgical blade, three 1.5cm plugs from the choice areas of the plate were separated as follows: OP50 choice, PA14 choice and Indecision choice (point of insert) and plated onto respective food plates. After 24 hours, the plates were sexed to determine the ratio of each sex in the choice made.

### Fast-Killing assay

The fast killing plates were prepared as described^25^. Briefly, PSG media (Peptone-Glucose-Sorbitol (1% Bacto-Peptone; 1% NaCl; 1% glucose; 0.15M sorbitol; 1.7% Bacto-Agar)) was poured onto 3cm plates and then left to dry. The bacterial cultures were prepared as described above. 10 μl of the overnight culture of PA14 was then used to inoculate the plates and spread evenly using a spreader. The worms were also prepared as described above; sex-separated young adults of each sex (50-80 animals) were introduced to the killing plates and incubated at RT for 21 hours; the number of live worms was counted every 6 hours, with dead worms removed from the plates. Once tabulated, the survival percentage was calculated. The assay was done in three technical replicates.

### Slow-killing assay

The slow-killing assay was based on previous methods^25^ with slight modifications. The bacterial cultures and plates were prepared as described above, except 20 μl of the overnight culture of PA14 was used to inoculate regular NGM plates and spread evenly using a spreader. Before introducing the worms, 40 μl of 100 mM FUDR was dropped on to the sides of the plate. The worms were prepared as described above, and sex-separated young adults of each sex (40-100 animals) were introduced to the killing plates and incubated at RT for 96 hours, during which, the number of live worms was counted every 24 hours, with dead worms removed from the plates. Once tabulated, the survival percentage was calculated. The assay was done in two biological replicates.

### Lawn-leaving assay

The bacterial cultures and plates were prepared as described above, except 30 μl of the overnight culture of OP50 or PA14 inoculum used to inoculate normal 6 cm NGM plates. Next, sex-separated age-synchronized animals were introduced and left at RT for 6-8 hours, and then the number of worms inside or outside the lawn was counted.

### Chemotaxis Assay

Olfactory preference assays for Diacetyl and Pyrazine was conducted as described in ^24,40,102^, with slight modifications. Animals were sex-separated at late L4s, washed and placed on a 10 cm plate equidistant from two spots of ethanol-diluted odors. After 60 mins, the animals within 1.5 cm of each odor spot was manually counted to achieve the olfactory preference index, OPI = (b-a)/(a+b), where a and b are the two odor spots.

### Sex-exposed learning assay

The assay for the virgin groups was carried out as described above. However, for the mated groups, hermaphrodites were introduced into the training plates at the start of the assay, and at the end of 6 hours, the males were separated, and the downstream processing was continued as usual.

### IRG-1 imaging

Age-synchronized animals were incubated on OP50 or PA14 training plates for 6 hours and imaged under a Zeiss LSM 880 confocal microscope at 25x magnification on an agar pad. The PA14 groups were imaged first and the OP50 groups within 8 hours of incubation. Each image file was subjected to the same post processing by converting it to 8-bit image z-projected using maximum intensity and then subtracting background by rolling ball radius of 50 pixels and thresholded between 4-5% before selecting the Region of Interest, which ran from start of the gut until the end of the anterior region, using FIJI and the mean intensity was quantified.

### AIY Rab-3 mCherry imaging

Age-synchronized animals were incubated on OP50 or PA14 training plates for 6 hours and imaged under a Zeiss LSM 880 confocal at 63x magnification on an agar pad. Each image file was subjected to the same post processing by converting it to 8-bit image, z-projected using maximum intensity and then subtracting background using rolling ball radius of 50 and thresholded to generate appropriate mask using analyze particles before selecting the Region of Interest in the red channel for mean intensity quantification. Zone 2 and Zone 3 was identified as previously described ^103^ and the mean intensity was quantified. To determine the delta between the training states, the males were normalized to the hermaphrodites, with hermaphrodite mean intensity taking a value of one.

### Calcium imaging

Overnight cultures of OP50 and PA14 were centrifuged at 4000g for 10 minutes and the supernatant was then filtered using a 0.22-micron syringe filter. Age-synchronized animals after 6 hours of training were used for imaging. Calcium imaging was performed according to Pechuk et al. ^8^, with changes made to the stimulus type and duration of stimulus. Briefly, the imaging chips were fabricated according to Chronis et al.^103^, with the help of the Nanofabrication Unit at the Weizmann Institute of Science. Two pumps control the flow of OP50 supernatant and PA14 supernatant into the microfluidic chip. The solutions were channeled through PVC tubes and using a manual switch, the stimulus was presented to the worm, through stainless steel connectors. The pumping rate during experiments was ∼0.005 ml/min. The worms were loaded into worm inlet of the chip by using a 1 ml syringe that had worms suspended in OP50 supernatant. 2 min habituation was given prior to imaging with the lasers on. To ensure smooth imaging, 10 mM Levamisole was added to all supernatants and 50 μM rhodamine B was added only to the PA14 supernatant to visualize stimulus delivery. Confocal imaging was done using a 40x magnification water objective.

The total imaging duration was 40 sec, constituting 544 frames in total, and the stimulus duration was 12 sec for AWC, 11 sec for AWB, and 15 sec for AIY, after an equal amount of initial time to capture baseline for each neuron respectively. All imaging files were exported as tiff files and the GCaMP fluorescence intensity was analyzed using FIJI. ROIs (regions of interest) of the somas or the axon were drawn manually to best represent the signal, and their mean grey values were quantified (Source Data). Downstream data processing was performed using MATLAB; for each worm, the baseline fluorescent level (F0) was calculated by averaging the mean grey values of 100 frames before stimulus delivery, then for each frame, the ΔF was calculated by subtracting F0 from the value of that time point, and the result was divided by F0, to normalize differences in the fluorescence baseline levels between individuals (ΔF/F0). A similar analysis was carried out in the stimulus channel of the images to align the responses to the start of the stimulus (AIY) or the end of the stimulus (AWB and AWC). Once aligned, the ON response was defined as the difference in mean of 10 seconds post stimulus start and the mean of 5 seconds pre-stimulus start. OFF response was defined as the difference in mean of 10 seconds post stimulus stop and the mean of 5 seconds pre-stimulus start to keep the baseline for both ON and OFF responses consistent. The latency of the response was defined as the time it took to reach 2 times the SD of the mean before stimulus start. A similar threshold was used to calculate the time of neuronal activity from the start of the stimulus to the end of the recording for each animal.

For calcium imaging with 2-nonaone as the stimulus, the same protocol was followed as described above, except that the buffer used was NGM buffer and the stimulus was 10^5^ diluted 2-nonaone.

### Transcriptomics analysis

Approximately 700–1000 animals of each sex were subjected to PA14 or OP50 training for 6 hours and washed with M9 to remove any bacterial remnants. Each training group for each sex had 4 biological replicates. The worm pellet was flash-frozen in 500μl TRIzol using liquid nitrogen and stored at −80°C. Total RNA was extracted using the TRIzol LS (Invitrogen) protocol. The RNA from the post isopropanol precipitation step was resuspended in extraction buffer from the PicoPure (Arcturus) RNA isolation kit, and further processing was performed according to the manufacturer’s recommendations. After determining the RNA concentration, using a Qubit Fluorometer (model number 4, Invitrogen), and quality, using a Bioanalyzer (Agilent), the samples with a RIN-Score >8 (Supplementary Data 1) were used to create RNA-seq libraries, as previously described using the MARS-seq procedure^49^. The FASTQ files were then processed using the user-friendly transcriptome analysis pipeline (UTAP) (https://utap.wexac.weizmann.ac.il/)^104^. The reads were aligned against the *C. elegans* reference genome [hWBcel235](https://www.ncbi.nlm.nih.gov/datasets/genome/GCF_000002985.6/). The data thus captured was analyzed against the Wormbase database for Gene IDs. Using Python, PCA plots were plotted using seaborn. The data was filtered by sex using the following parameters: Basemean >5 and p-value >0.05 from each sex (naïve vs trained) for all analyses and genes with Basemean >5 was taken ahead for only volcano plots. The p-values for neuronal genes were corrected using the Benjamini-Hochberg method (Supplementary Data 2). The heatmaps and volcano plots were plotted using the Heatmap and EnhancedVolcano functions of R studio (2022.12.0+353), respectively. The resultant data was further taken for GO term analysis of genes from each sex (naïve vs trained) using ShinyGo^105^ and the circular barplot was made using ggplot of R studio (2022.12.0+353).

### RT-PCR

Total RNA was extracted for all groups using the protocol described above in ‘Transcriptomics analysis’. The isolated RNA was converted to cDNA using random hexamers from the SuperScript^TM^ IV First-Strand Synthesis system. After the concentration was assessed using NanoDrop, qPCR reactions were set up in 10ul reaction mixtures as per manufacturer instructions using Fast SYBR Green Master Mix and the StepOnePlus^TM^ Real-Time PCR system. Ct values were used to quantify the relative expression of the target gene using the 2 − ΔΔCt method. *nhr-23* was used as the housekeeping gene.

### Dye staining of *npr-5::GFP* animals

Animals at the YA stage were washed with M9 buffer and incubated in 1ml M9 and 5μl DiD dye (Vybrant DiD Cell-labelling solution, ThermoFisher), for 1 hour, on a rotor at ∼40 rpm. The worms were then transferred to food plates and the M9 allowed to dry. The animals were then sex-separated and transferred to either OP50 or PA14 training plates and incubated for 6 hours. Next, the worms were transferred to an agar pad and anesthetized with 5μl of 100mM NaN3 and imaged under a Zeiss LSM 880 using a 40x water objective. The Region of Interest was selected in FIJI, using the DiD staining, for the positions of ASJ and AWB, while a co-injection marker was used to assess the position of AIY. The mean intensity was quantified after background correction and thresholding.

### CRISPR/Cas-9 mediated *npr-5* deletion

The CRISPR strategy was based on Paix et al.^106^. Recombinant cas9 (IDT) was injected into N2 gonads with tracrRNA (IDT) and three crRNAs, one targeting the dpy-10 locus (Paix et al. 2015) and two flanking the *npr-5* locus (GATAACGTTGCTGATGGTGG, located 52bp downstream of *npr-5* start codon, and GAGACGGGCAAGATGAACAC, located 54bp downstream of stop codon). The repair template was the ssODN CCATTTCGACCATTTCAACCACAACGACTCCCTCCGTTCATCTTGCCCGTCTCTCGAAAAATTCCATTAT, composed of 35nt homology arms from each side of the *npr-5* locus. Rol/dpy F1 progeny were screened by PCR to identify the *npr-5(ety11)* deletion allele. Three primers were included in the reaction: TTCGATGCATGGTTAGTTCG (*npr-5* CR_F1), TTTCCATGTCACTCGTCAGC (*npr-5* CR_R1), GGAAATTCCCGGAAAATGAT (*npr-5* CR_R2), resulting in a 347bp band in WT animals and a 577bp band in ety11 animals. The *syb5786* allele was generated by SunyBiotech.

### Transgenic strains/molecular cloning

The feminization of the gut was generated by expressing *tra-2[intracellular]* under the *elt-2* promoter, cloned by Gibson assembly^90^. *elt-2* promoter was amplified from a plasmid provided by Prof. Sivan Korenblit.

To express *npr-5* cDNA specifically in AWB, *npr-5* cDNA was obtained from Genescript and fused to 3.5kb promoter of *str-1* and upstream of SL2::NLS::tagRFP::unc-54 3’UTR using PCR fusion^107^.

Primers used are in Supplementary Data 3.

### RNAi

RNA interference of select genes was induced using the feeding method ^108^. Briefly, L4 hermaphrodites were fed HT115 bacteria carrying the dsRNA for the relevant genes including a control empty vector. Multiple plates for each gene were maintained to achieve sufficient young adult males to perform the learning assay as described above. Fresh *pos-1* control was prepared for each RNAi experiment to verify efficiency (Only batches > 95% were used for behavioral assays).

### Quantification and statistical analysis

Each figure legend contains details of the quantification, statistical tests performed and the sample sizes of each individual experiment. All statistical analysis was performed on GraphPad Prism 10.

## Acknowledgments

We thank the members of Oren-Suissa lab for their insights regarding the manuscript. We especially thank Dr. Yehuda Salzberg who helped us in generating the *npr-5* CRISPR strain used in this study. We would like to thank Mario De Bono for furnishing us with much-needed strains. Some strains were provided by the CGC, which is funded by the NIH Office of Research Infrastructure Programs (P40 OD010440). We would also like to thank Prof. Sivan Korenblit for plasmids and Prof. Ravid Straussman for providing us with *S. aureus*. We would like to extend our gratitude to the Nano-Fabrication unit and the Genomics Sandbox unit at the Life Science Core Facility of Weizmann Institute of Science for helping us with the PDMS chips and the transcriptomics, respectively. We thank WormBase, the online database of *C. elegans* genes, for the excellent curation of various information regarding each gene. MOS acknowledges financial support from the European Research Council ERC-2019-STG 850784, Israel Science Foundation grant 961/21, Dr. Barry Sherman Institute for Medicinal Chemistry, Sagol Weizmann-MIT Bridge Program and the Azrieli Foundation. MOS is the incumbent of the Jenna and Julia Birnbach Family Career Development Chair.

## Authors contribution

SPK conducted and analyzed the experiments. RH performed the library preparation and sequencing for the transcriptomics experiment. AG performed and analyzed the *npr-5* dye staining experiment. MOS supervised and designed the experiments. SPK and MOS wrote the paper.

## Data availability

All data generated and used for analysis are included in the manuscript along with the supplementary information files and Source Data files. The sequence data generated in the study has been deposited in the GEO database under the accession code [hGSE273091](https://www.ncbi.nlm.nih.gov/geo/query/acc.cgi?acc=GSE273091). The RNA-seq data is provided in Supplementary Data 1 and Supplementary Data 2. Quantifications for calcium imaging are added to the Source Data. Raw images for all the imaging experiments can be found at (<u>10.34933/2c05a150-4238-4273-93ec-b419d2c5429f</u>). Source Data are provided in this paper.

## Code availability

Code generated in this study is available at (10.34933/2c05a150-4238-4273-93ec-b419d2c5429f).

