## Supplemental File for "Modulation by NPYR underlies experience-dependent, sexually dimorphic learning"

### Supplementary Material

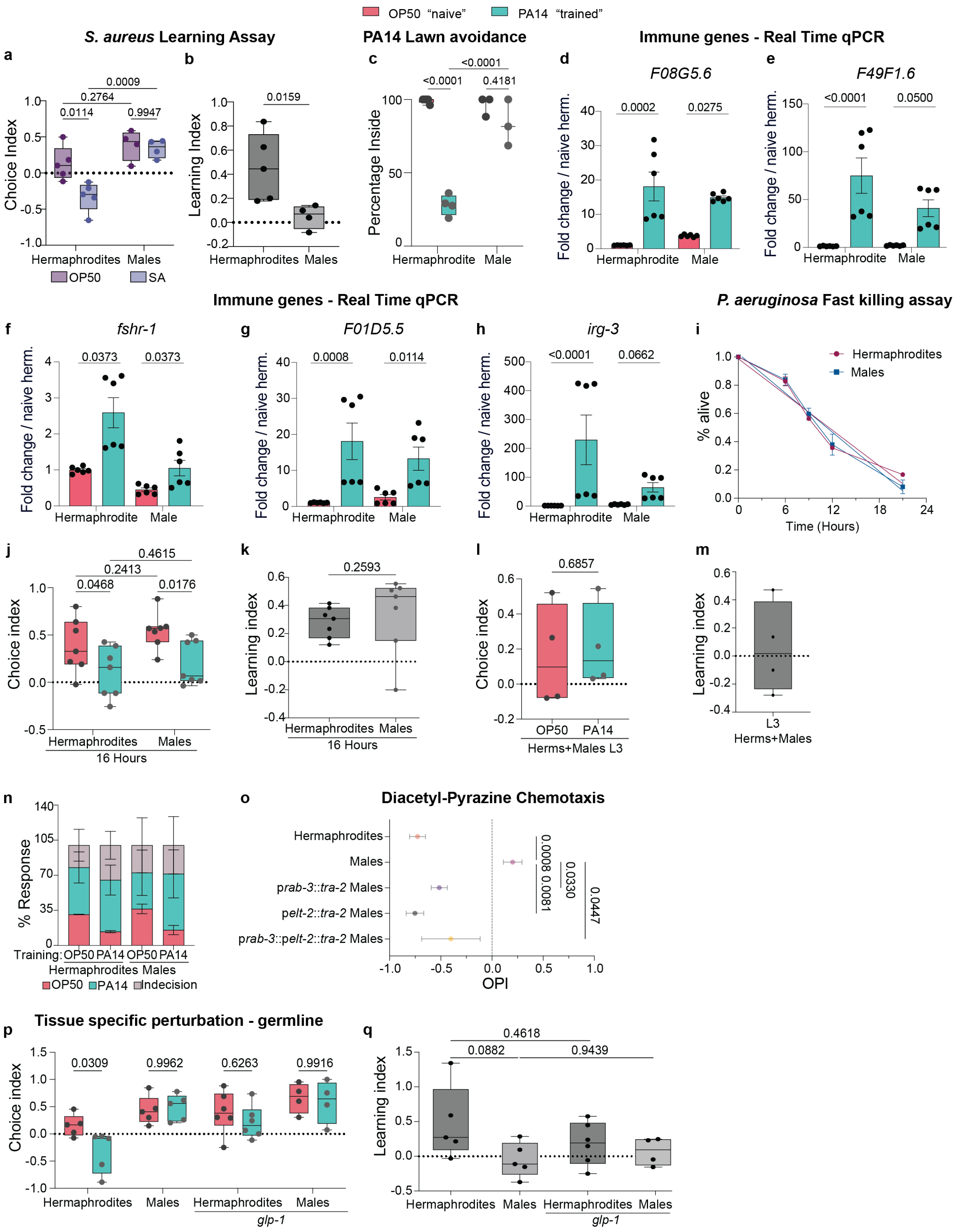


2

3 **Supplementary Fig. 1. PA14 short-term learning is stage-, tissue- and sex-specific.**

**a** The Choice Index of hermaphrodites (OP50, PA14: N = 5) and males (OP50, PA14: N = 5) subjected to training paradigm using *S. aureus*. **b** The learning indices of hermaphrodites and males subjected to *S. aureus* training. **c** Percentage of hermaphrodites (OP50 N = 5, PA14 N = 4) and males (OP50, PA14 N = 3) inside the lawn on either OP50 (magenta) or PA14 (cyan) for 6-8 hours. **d-h** Real-time qPCR analysis of the immune response genes *F08G5.6*, *F49F1.6*, *fshr-1*, *F01D5.5* and *irg-3* mRNA normalized to naïve hermaphrodite expression (N = 2 biological samples with 400-600 animals per group). **i** Percentage of live hermaphrodites and males in fast-killing assay. **j-k** Choice **(j)** and Learning **(k)** indices of hermaphrodites (OP50, PA14 N = 7) and males (OP50, PA14 N = 7) after 16 hours of exposure to PA14 (cyan) or OP50 (magenta). **(l-n)** Choice **(l)** and Learning **(m)** indices and response percentage **(n)** of L3 hermaphrodites (OP50, PA14 N = 4) and males (OP50, PA14 N = 4). **o** Diacetyl-pyrazine olfactory preference assay in wild-type and transgenic animals. '+1' indicates strong preference for pyrazine and '-1' indicates strong preference for diacetyl. Wild-type hermaphrodites and males (N = 8), pan-neuronally feminized (p*rab-3*::*tra-2(ic)*) males (N = 9), gut-feminized (p*elt-2*::*tra-2(ic)*) males (N = 4), and pan-neuronally and gut feminized (p*rab-3*::*tra-2(ic)*;p*elt-2*::*tra-2*) males (N = 6). OPI, Olfactory Preference Index (see methods). **p-q** Choice **(p)** and Learning **(q)** indices of wild-type and germline mutant hermaphrodites and males, showing no learning effect in males (wild-type hermaphrodites and males: OP50, PA14 N = 5; *glp-1* hermaphrodites: OP50, PA14 N= 6; *glp-1* males: OP50, PA14 N = 6). In all supplemental figures, n represents the number of animals tested, and N represents the number of biological repeats. Statistical tests: In **(a, c, i, j, p)**, a two-way ANOVA with Šídák's multiple comparisons test. In **(b, k, l)**, a Mann Whitney test. In **(d-h)** Kruskal-Wallis test with Uncorrected Dunn's multiple comparisons test. In **(n)**, a two-way ANOVA with Tukey's multiple comparisons test. In **(o)**, Kruskal-Wallis test with Dunn's multiple comparisons test. In **(q)**, a One-Way ANOVA with Šídák's multiple comparisons test. Graphs are Box and Whisker plots with mean and entire data range. P values are indicated in the figure.

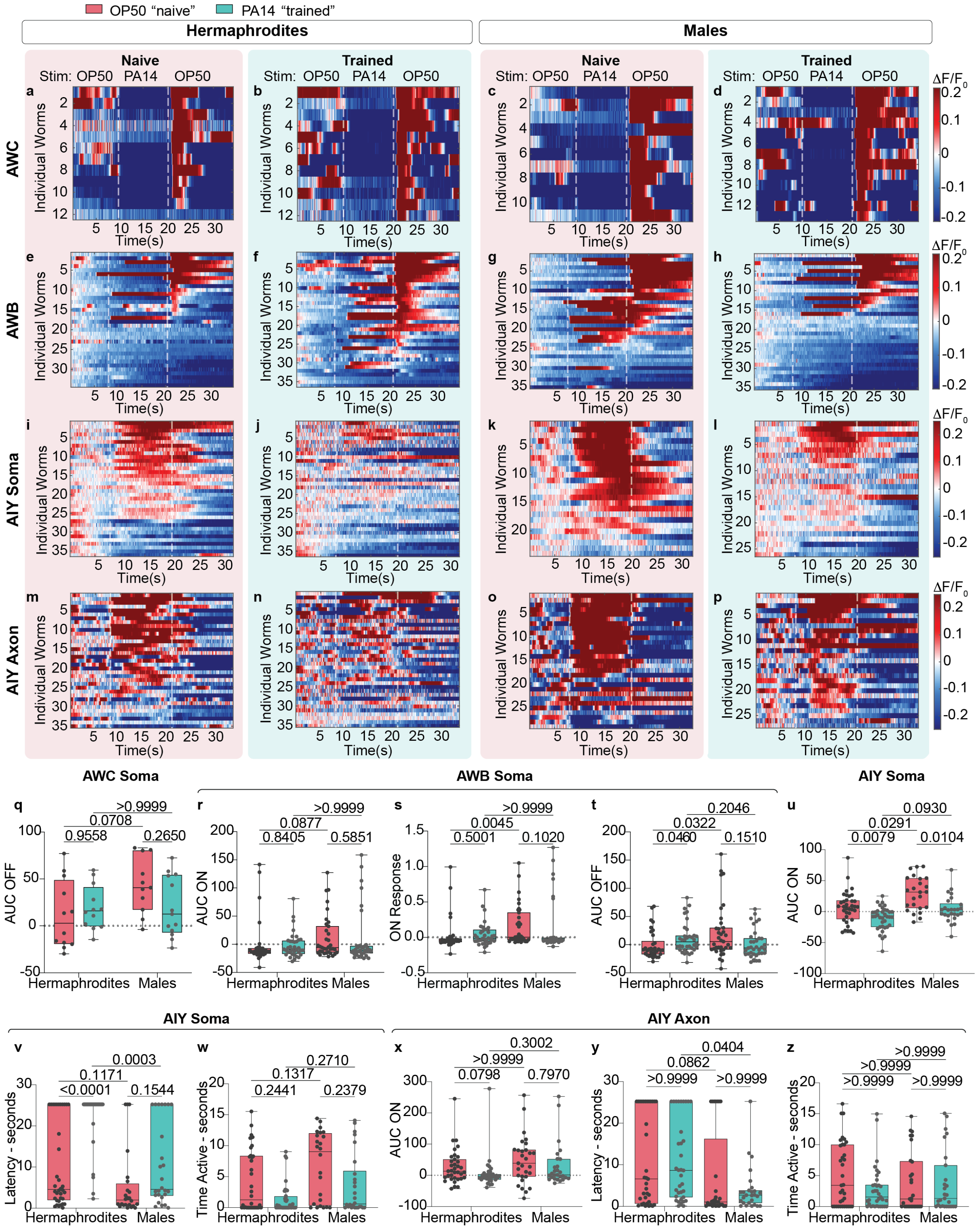


**Supplementary Fig. 2. Distinct dimorphic information processing towards PA14 stimuli.**

**a-p** From left to right, heatmap of GCaMP signals of select neurons in naïve (magenta) and trained (cyan) hermaphrodites and males. **(a-b)** AWC calcium levels in naive (n = 12) and trained (n =

12) hermaphrodites. **(c-d)** AWC calcium levels in naive (n = 11) and trained (n = 13) males. **(e-f)** AWB calcium levels in naive (n = 33) and trained (n = 35) hermaphrodites. **(g-h)** AWB calcium levels in naive (n = 35) and trained (n = 35) males. **(i-j)** AIY soma calcium levels in naïve (n = 34) and trained (n = 36) hermaphrodites. **(k-l)** AIY soma calcium levels in naïve (n = 24) and trained (n = 26) males. **(m-n)** AIY axon calcium levels in naïve (n = 35) and trained (n = 35) hermaphrodites. **(o-p)** AIY axon calcium levels in naïve (n = 27) and trained (n = 26) males. **q** AUC of the OFF response of AWC in both sexes of naïve and trained animals. **r-t** Quantification of **(r)** AUC, of the **(s)** ON response and **(t)** AUC of OFF response of AWB of both sexes of naïve and trained animals. **u-w** Quantification of the **(u)** AUC of ON response of AIY soma and the **(v)** latency of ON response and **(w)** duration of ON response of both sexes in naïve and trained animals. **x-z** Quantification of **(x)** AUC of ON response of AIY axon and the **(y)** latency of ON response and **(z)** duration of ON response of both sexes in naïve and trained animals. Statistical tests: in **(q)** a two-way ANOVA with Šídák's multiple comparisons test. In **(r-z)**, a Kruskal-Wallis test with Dunn's multiple comparisons test. Box and Whisker plots with mean and entire data range. P values are indicated in the figure.

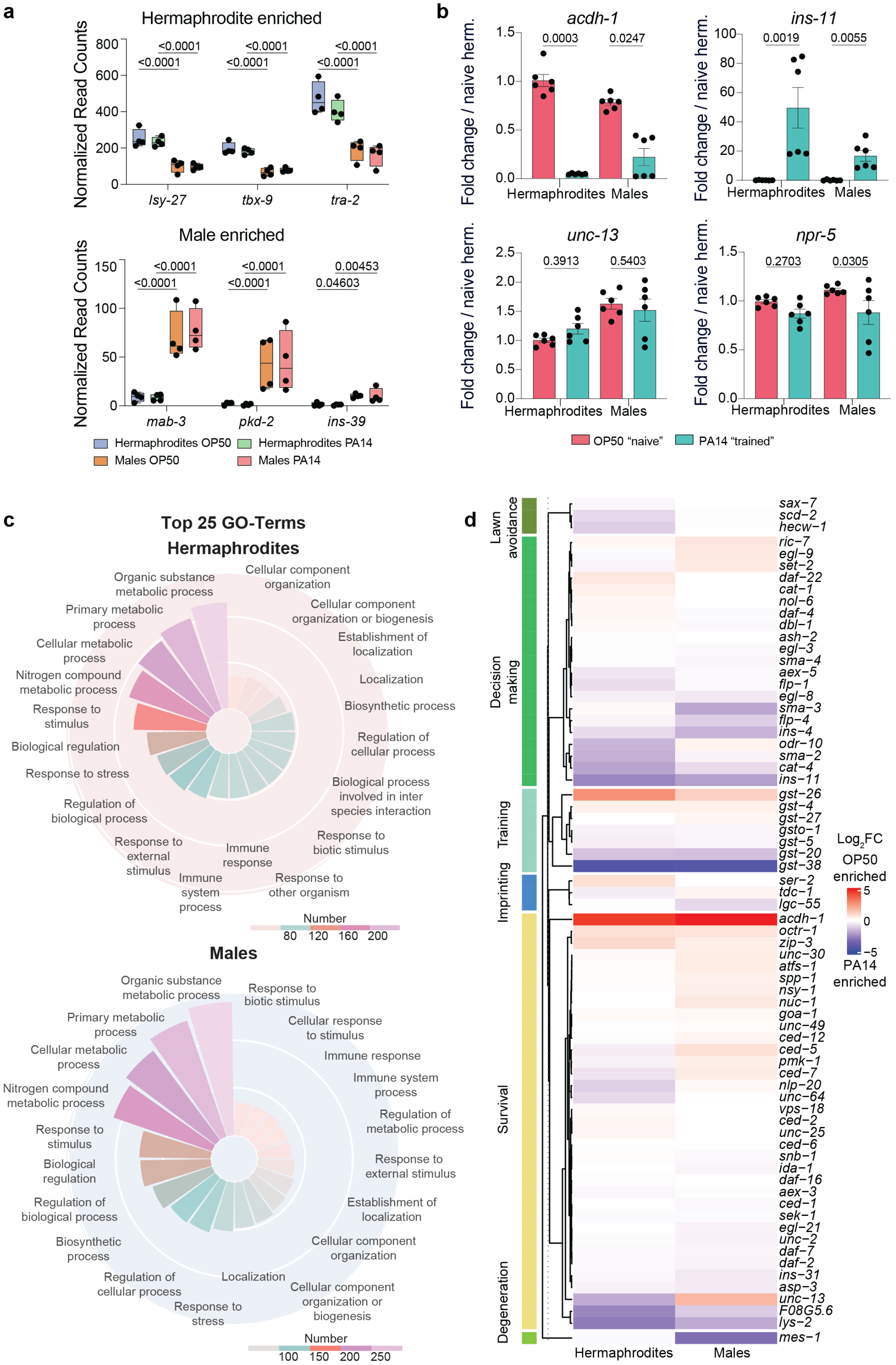
a
Hermaphrodite enriched
Normalized Read Counts
800
600
400
200
0
<0.0001
<0.0001
<0.0001
<0.0001
<0.0001
<0.0001
lsy-27
tbx-9
tra-2
Male enriched
Normalized Read Counts
150
100
50
0
<0.0001
<0.0001
<0.0001
<0.0001
0.00453
0.04603
mab-3
pkd-2
ins-39
Hermaphrodites OP50
Hermaphrodites PA14
Males OP50
Males PA14
b
acdh-1
Fold change / naive herm.
1.5
1.0
0.5
0.0
0.0003
0.0247
Hermaphrodites
Males
ins-11
Fold change / naive herm.
100
80
60
40
20
0
0.0019
0.0055
Hermaphrodites
Males
unc-13
Fold change / naive herm.
2.5
2.0
1.5
1.0
0.5
0.0
0.3913
0.5403
Hermaphrodites
Males
npr-5
Fold change / naive herm.
1.5
1.0
0.5
0.0
0.2703
0.0305
Hermaphrodites
Males
OP50 "naive"
PA14 "trained"
c
Top 25 GO-Terms
Hermaphrodites
Organic substance metabolic process
Cellular component organization
Primary metabolic process
Cellular component organization or biogenesis
Cellular metabolic process
Establishment of localization
Nitrogen compound metabolic process
Localization
Response to stimulus
Biosynthetic process
Biological regulation
Regulation of cellular process
Response to stress
Biological process involved in inter species interaction
Regulation of biological process
Response to biotic stimulus
Response to external stimulus
Immune response
Immune system process
Response to other organism
Number
80
120
160
200
Males
Organic substance metabolic process
Response to biotic stimulus
Primary metabolic process
Cellular response to stimulus
Cellular metabolic process
Immune response
Immune system process
Nitrogen compound metabolic process
Regulation of metabolic process
Response to stimulus
Biological regulation
Response to external stimulus
Regulation of biological process
Establishment of localization
Biosynthetic process
Cellular component organization
Regulation of cellular process
Localization
Cellular component organization or biogenesis
Response to stress
Number
100
150
200
250
d
Lawn avoidance
Decision making
Training
Imprinting
Survival
Degeneration
sax−7
scd−2
hecw−1
ric−7
egl−9
set−2
daf−22
cat−1
nol−6
daf−4
dbl−1
ash−2
egl−3
sma−4
aex−5
flp−1
egl−8
sma−3
flp−4
ins−4
odr−10
sma−2
cat−4
ins−11
gst−26
gst−4
gst−27
gsto−1
gst−5
gst−20
gst−38
ser−2
tdc−1
lgc−55
acdh−1
octr−1
zip−3
unc−30
atfs−1
spp−1
nsy−1
nuc−1
goa−1
unc−49
ced−12
ced−5
pmk−1
ced−7
nlp−20
unc−64
vps−18
ced−2
unc−25
ced−6
snb−1
ida−1
daf−16
aex−3
ced−1
sek−1
egl−21
unc−2
daf−7
daf−2
ins−31
asp−3
unc−13
F08G5.6
lys−2
mes−1
Hermaphrodites
Males
Log2FC
OP50 enriched
5
0
−5
PA14 enriched

**Supplementary Fig. 3. Non-dimorphic macro molecular landscape.**

**a** Normalized RNA-seq expression values of hermaphrodite enriched (*lsy-27, tbx-9, tra-2*) and male enriched (*mab-3, pkd-2, ins-39*) genes indicating the sex-specificity of the transcriptomic dataset. **b** Real-time qPCR analysis of the select genes *acdh-1*, *ins-11*, *unc-13* and *npr-5* mRNA normalized to naïve hermaphrodite expression (N = 2 biological samples with 400-600 animals per group). **c** GO term analysis of naïve vs. trained hermaphrodites and males, showing similar macro transcriptomic changes. **d** Heatmap of $log_2$ fold-change of previously described PA14-relevant genes in naïve vs. trained hermaphrodites and males from the whole transcriptomics dataset. Statistical tests: in **(a)**, Two-sided Wald's test and in **(b)** Kruskal-Wallis test with Uncorrected Dunn's multiple comparisons test. Bar graph with SEM and individual data points. P values are indicated in the figure.

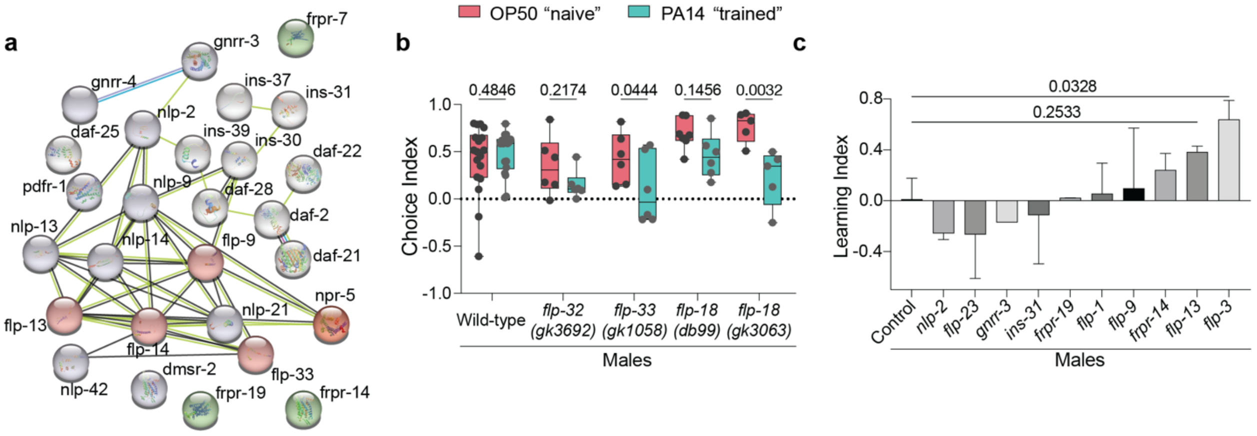


**Supplementary Fig. 4. Neuropeptidergic network modulates male learning**

**a** Schematic of all the neuropeptide-related genes from our dataset using String Protein Ensemble[1] showing interactions between several candidate genes, including *npr-5*. **b** Choice Indices of wild-type (OP50 N = 18, PA14 N = 18), *flp-32 (gk3692)* mutant (OP50, PA14, N = 6), *flp-33 (gk1058)* mutant (OP50, PA14 N = 6), *flp-18 (db99)* mutant (OP50 N = 7, PA14 N = 6) and *flp-18 (gk3063)* mutant (OP50, PA14 N = 5) males showing peptidergic regulation of male learning. **c** Learning indices of RNAi-treated males showing *flp-3* as a likely neuromodulator (control: N = 5; *gnrr-3*: N = 1; *nlp-2*, *flp-23*, *ins-31*, *frpr-19*, *flp-1*, *flp-9*, *frpr-14*, *flp-13*, *flp-3*: N = 2). Statistical tests: in **(b)**, a two-way ANOVA with Tukey's multiple comparisons test. In **(c)**, a Kruskal-Wallis test with Dunn's multiple comparisons test. Box and Whisker plots with mean and entire data range. P values are indicated in the figure.

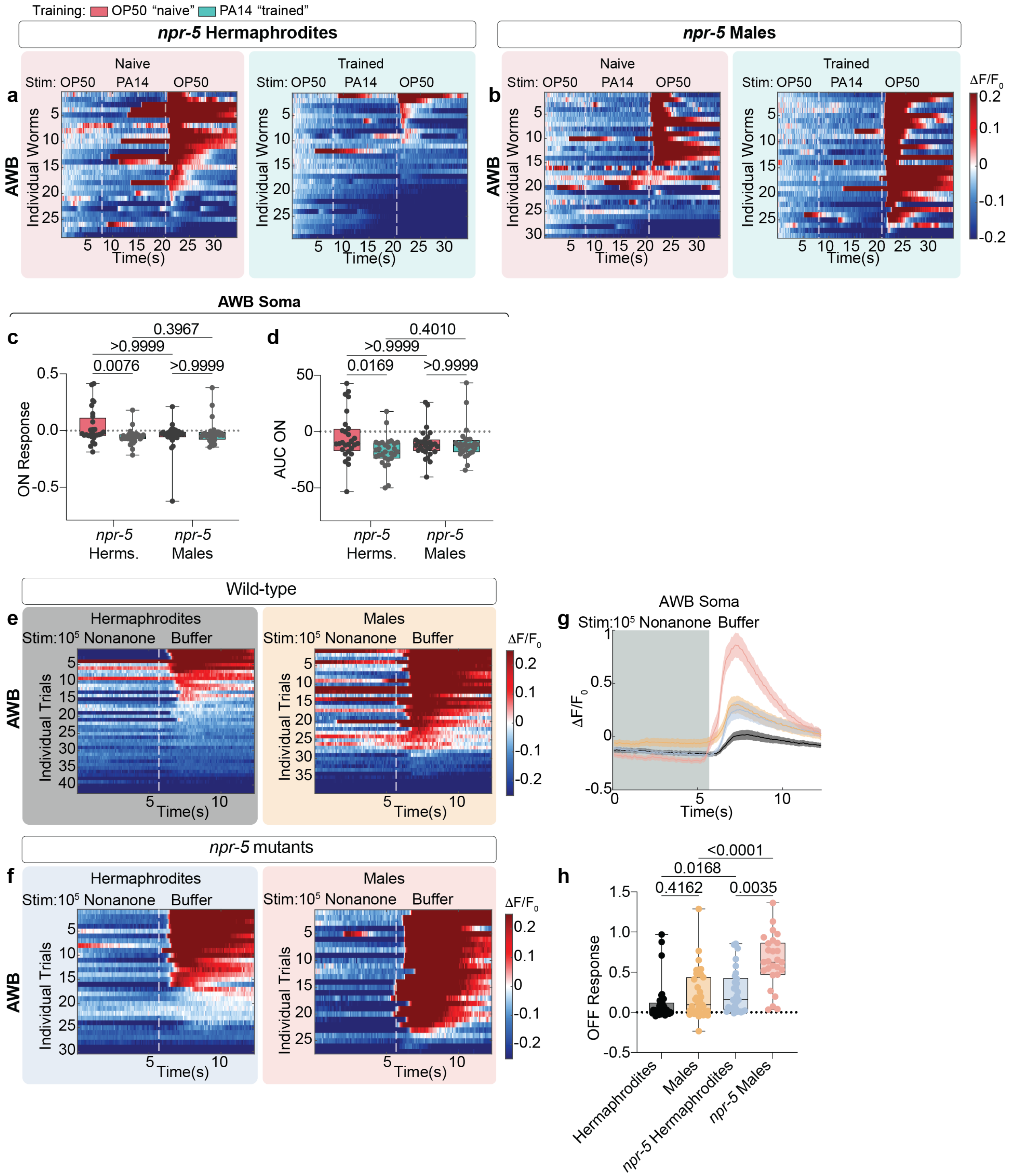


**Supplementary Fig. 5. NPR-5 alters AWB neuronal responses**

**a-b** From left to right, heatmap of GCaMP signals of AWB in naïve (magenta) and trained (cyan) *npr-5* mutant hermaphrodites and males. **(a)** AWB soma calcium response of naïve and trained *npr-5* mutant hermaphrodites (OP50 n = 28, PA14 n = 29) and **(b)** males (OP50 n = 30, PA14 n = 28) to PA14 supernatant exhibiting a reduction in the calcium response after the PA14 stimulus.

**c-d** Quantification of the **(c)** ON response and **(d)** AUC of OFF response of AWB of both sexes of naïve and trained *npr-5* mutant animals. **e-f** From left to right, heatmap of GCaMP signals of the soma of AWB neurons in **(e)** wild-type hermaphrodite (black; 42 trials from n = 14) and wild-type male (yellow; 39 trials from n = 13) and **(f)** *npr-5* hermaphrodite (blue; 30 trials from n = 10) and *npr-5* male (salmon; 27 trials from n = 9) to $10^5$ diluted 2-nonanone. **g** The SEM traces of calcium responses of both genotypes and sexes. The shaded region in the SEM traces corresponds to the exposure window of $10^5$ diluted 2-nonanone. **h** Quantification of OFF responses of both genotypes and sexes. In **(c-d, h)**, the statistical test used is Kruskal-Wallis with Dunn's multiple comparisons test. Box and Whisker plots with mean and entire data range. P values are indicated in the figure.

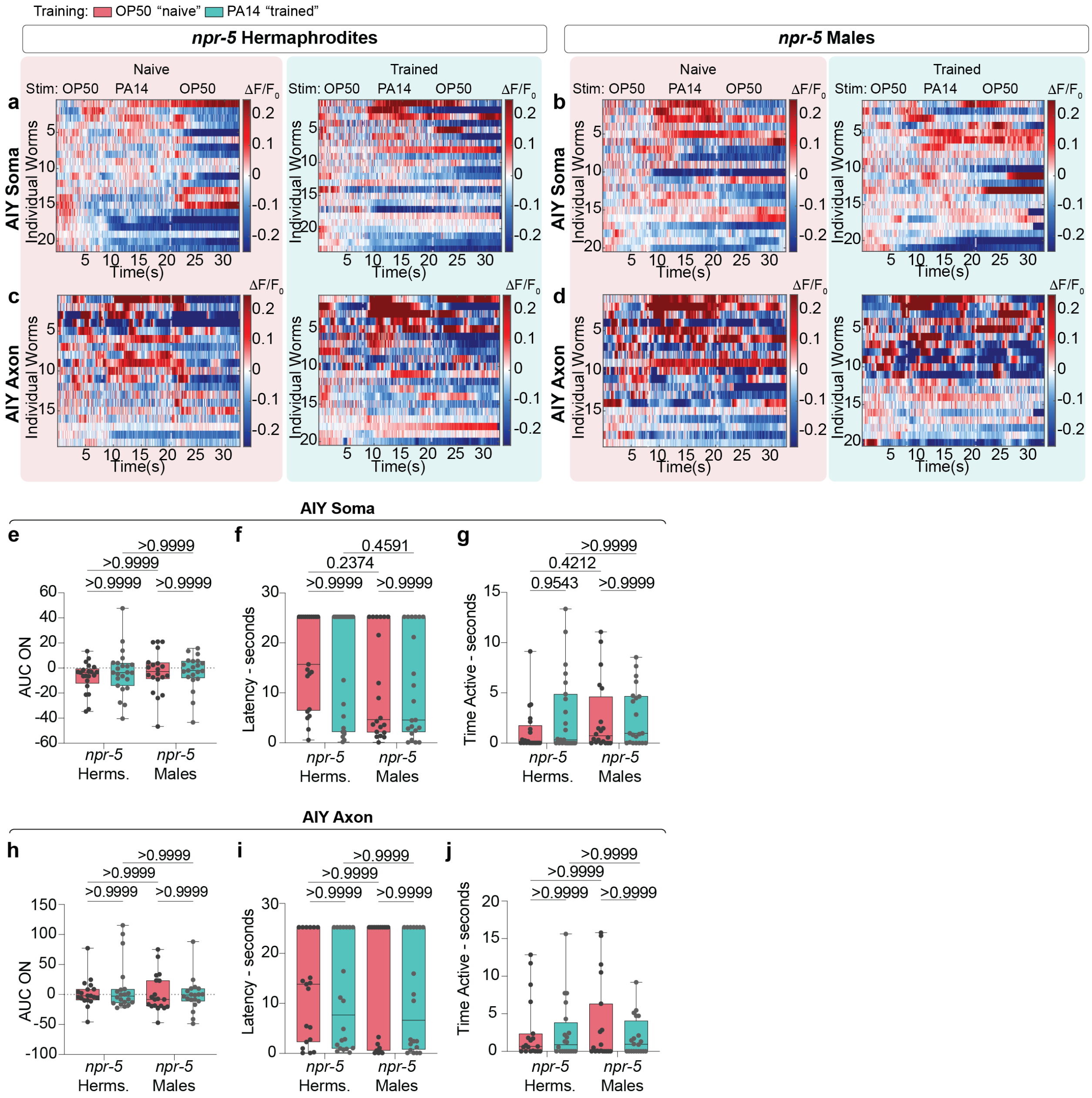


**Supplementary Fig. 6. NPR-5 is important for AIY activation states**

**a-d** From left to right, heatmap of GCaMP signals of AIY in naïve (magenta) and trained (cyan) hermaphrodites and males. **(a)** AIY soma calcium levels in naïve (n = 21) and trained (n = 23) *npr-5* mutant hermaphrodites. **(b)** AIY soma calcium levels in naïve (n = 20) and trained (n = 21) *npr-5* mutant males. **(c)** AIY axon calcium levels in naïve (n = 20) and trained (n = 19) *npr-5* mutant hermaphrodites. **(d)** AIY axon calcium levels in naïve (n = 19) and trained (n = 20) *npr-5* mutant males. **e-g** Quantification of the **(e)** AUC of ON response, **(f)** latency of ON response and **(g)** duration of ON response of both sexes in AIY soma of naïve and trained *npr-5* mutant animals. **h-j** Quantification of the **(h)** AUC of ON response, **(i)** latency of ON response and **(j)** duration of

ON response of both sexes in AIY axon of naïve and trained *npr-5* mutant animals. In **(e-j)**, the statistical test used is Kruskal-Wallis with Dunn's multiple comparisons test. Box and Whisker plots with mean and entire data range. P values are indicated in the figure.

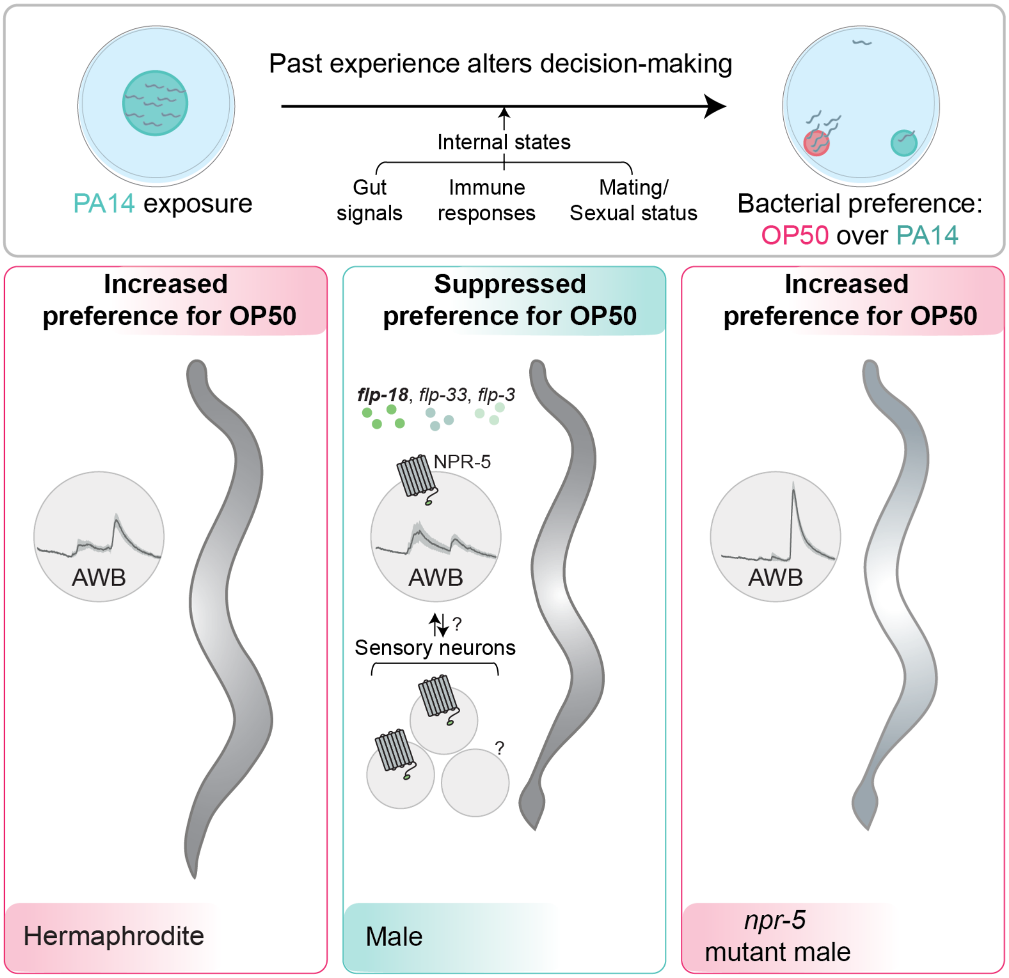


**Supplementary Fig. 7. Model for male PA14 aversive learning**

Schematic of the sexually dimorphic response to PA14. Short-term exposure to PA14 triggers changes in internal states, causing hermaphrodites to prefer OP50 to PA14, represented as a higher OFF response in AWB neurons. In males, however, this choice is suppressed. AWB shows reduced OFF response due to *npr-5,* and the behavior is modulated by several neuropeptides, including *flp-18* and additional regulators, possibly form the gut and other neurons. In *npr-5* mutant males, there is a heightened AWB OFF response, similar to wild-type hermaphrodites and subsequently preference to OP50 after exposure.

**Supplementary Data 1. (separate file)**

Raw read counts, RINe scores, raw and normalized read counts per gene, per sample for both naïve and trained hermaphrodites and males.

**Supplementary Data 2. (separate file)**

Neuro-relevant genes, genes implicated in literature along with GO term analysis for male- and hermaphrodite-enriched samples after training.

**Supplementary Data 3. (separate file)**

List of strains, reagents and resources used in this study.
